# Mutations in a single signaling pathway allow growth on a different solvent than water

**DOI:** 10.1101/692889

**Authors:** Caroline Kampmeyer, Jens V. Johansen, Christian Holmberg, Magnus Karlson, Sarah K. Gersing, Heloisa N. Bordallo, Birthe B. Kragelund, Mathilde H. Lerche, Isabelle Jourdain, Jakob R. Winther, Rasmus Hartmann-Petersen

## Abstract

Since life is completely dependent on water, it is difficult to gauge the impact of solvent change. To analyze the role of water as a solvent in biology, we replaced water with heavy water (D_2_O), and investigated the biological effects by a wide range of techniques, using the fission yeast *Schizosaccharomyces pombe* as model organism. We show that high concentrations of D_2_O lead to altered glucose metabolism, growth retardation, and inhibition of meiosis. However, mitosis and overall cell viability were only slightly affected. After prolonged incubation in D_2_O, cells displayed gross morphological changes, thickened cell walls as well as aberrant septa and cytoskeletal organization. RNA sequencing revealed that D_2_O causes a strong downregulation of most tRNAs and triggers activation of the general stress response pathway. Genetic screens identified several D_2_O sensitive mutants, while mutants compromised in the cell integrity pathway, including the protein kinase genes *pmk1*, *mkh1*, *pek1* and *pck2*, that control cell wall biogenesis, were more tolerant to D_2_O. We speculate that D_2_O affects the phospholipid membrane or cell wall glycans causing an activation of the cell integrity pathway. In conclusion, the effects of solvent replacement are pleiotropic but the D_2_O-triggered activation of the cell integrity pathway and subsequent increased deposition of cell wall material and septation problems appear most critical for the cell growth defects.

## Introduction

Although in principle life might be possible in other solvents such as ammonia or formamide [1], life, as we know it, is completely dependent on water. As solvent, water provides a liquid phase that facilitates chemical reactions by allowing reactants to encounter each other, but also ensures that various biomolecules, organelles and tissues can maintain a functional structure. Importantly, water also actively partakes in chemical reactions as a reactant (e.g. hydrolysis) or as a product (e.g. condensation). To some degree, organic solvents, such as DMSO, fulfill some of these requirements. However, these solvents are often toxic already at low concentrations [2] and do not generally engage in chemical reactions.

^1^H is by far the most abundant isotope of hydrogen and while ^2^H or deuterium (D) and D_2_O (heavy water) are fairly similar to ^1^H and light water, respectively, the chemical effects of isotope substitution is far stronger than seen for most other chemical elements [3]. Thus, when substituting water as solvent, there are very significant differences between light and heavy water. Both compounds are non-radioactive, but D_2_O is denser (1.1 vs. 1.0 g mL^-1^) and more viscous (1.25 vs. 1.00 mPa s at 20 °C) than H_2_O. It has a higher melting point (3.82 vs. 0 °C) as well as a higher boiling point (101.4 vs. 100 °C), consistent with the deuterium bonds in D_2_O being stronger than the corresponding hydrogen bonds in H_2_O. Accordingly, D_2_O is known to influence the conformational and functional properties of proteins and other macromolecules *in vitro* [4, 5].

Although the natural abundance of deuterium in water is less than 0.02%, it is interesting to study the cellular effects of D_2_O because its unique properties allow for a complete solvent replacement, and may therefore highlight the importance of the biophysical properties of H_2_O as solvent for biological processes. However, the cellular consequences of exchanging H_2_O with D_2_O have so far not been systematically addressed. When D_2_O replaces H_2_O in the cell, it affects reaction kinetics and the structure of macromolecules, giving rise to the so-called “solvent isotope effect” [5]. However, the deuterium atoms in D_2_O can also metabolically replace hydrogen atoms in biomolecules. When deuterium is metabolically incorporated into proteins and other biomolecules, the properties of C-D bonds in particular, which do not readily exchange with H, become important. Thus, C-D bonds may affect the structure and dynamics of macromolecules, resulting in the “deuterium isotope effect” [5]. Since hydrogen is much lighter than other biologically-relevant elements, the proportional isotope effects of deuterium are significantly stronger than e.g. those of ^13^C (vs. ^12^C) or ^18^O (vs. ^16^O) [3].

Soon after its discovery and large-scale manufacture, the first biological effects of D_2_O were described (for review see [5]). Today we know that small amounts of D_2_O are not toxic [6], and D_2_O can be used to measure the metabolic rate in humans [7, 8], or as a tracer for compliance in drug trials [9]. However, biological studies on D_2_O have shown that at high levels (>25% of body weight) D_2_O is toxic to animals and causes sterility [5]. In addition, heavy water has been reported to affect the period of circadian oscillations in *Drosophila* [10], while studies in rodents have shown that high D_2_O concentrations causes anemia and early death [11, 12]. In plant and animal cells, D_2_O inhibits mitosis [13, 14], which may be linked to increased microtubule stabilization [15, 16]. Bacteria can adapt to grow in 100% D_2_O [17–19], while budding yeast cells can tolerate up to 90% heavy water [20]. However, heavy water sensitivity has been described as a conditional phenotype in budding yeast and a D_2_O hypersensitive mutant in the ASP5 gene, encoding a cytosolic aspartate aminotransferase involved in nitrogen metabolism, has been isolated [20]. Recently, it was shown that the amount of deuterated metabolites decline in yeast cells during aging, and supplementing the growth medium with 50 % D_2_O increases the lifespan [21], presumably by slowing the metabolism. As a solvent, D_2_O has been reported to marginally increase the heat stability of macromolecules [4], including double stranded DNA [22, 23] and some proteins [24–26]. In addition, D_2_O can function as a chemical chaperone for misfolded proteins [27]. However, on the cellular level, D_2_O may lead to a decrease in heat tolerance [28].

Here, we analyzed the biochemical and physiological effects of D_2_O on the fission yeast *Schizosaccharomyces pombe*, a well characterized and genetically tractable eukaryotic model organism [29]. We found that the *solvent exchange* effect rather than the *isotope* effect caused a strongly reduced growth rate at high concentrations of D_2_O. Although the overall cell viability remained largely unaffected, D_2_O inhibited the glucose metabolism and caused gross morphological changes, thickened cell walls and abnormal cell septation. Transcriptomic analyses revealed that D_2_O triggered differential expression of numerous genes, including a strong reduction of tRNAs, and activation of the general stress response pathway. Genetic screens identified several D_2_O sensitive mutants, including a deletion in the *rga7* GTPase activating protein (GAP). Surprisingly, mutants in *pck2, mkh1*, *pek1,* and *pmk1*, that encode kinases of the cell integrity pathway, which controls cell wall biogenesis, were tolerant to the solvent exchange.

## Results

### Dramatically reduced growth of S. pombe exposed to high concentrations of D_2_O

The D_2_O was purchased as 99.8% D_2_O/0.2% H_2_O. For simplicity, when all regular deionized water in the growth medium was replaced with the 99.8% pure D_2_O we will in the following refer to these experiments as performed at 100% D_2_O, and similarly for experiments conducted at lower D_2_O concentrations. Importantly, the other components (e.g. glucose) in the media were, unless stated otherwise, not deuterated. Hence, even in our 100% D_2_O media some non-exchangeable protons are available to the cells and will be incorporated in biomolecules.

As a first step in our cell physiological analyses of D_2_O, we followed the growth of wild type S. pombe cells exposed to different concentrations of D_2_O. Pre-cultures, prepared in standard rich medium (YES medium with H_2_O), were washed and diluted with YES medium with 100%, 75%, 50%, 25% and 0% heavy water and growth was monitored by measuring the turbidity of the cultures. At D_2_O concentrations >50%, the cell growth rate was strongly reduced (Fig. 1AB) and the onset of growth inhibition occurred rapidly, already within the first generation (Fig. 1C). The effect was dosage dependent leading to an increase in doubling time from approximately 2.5 hours in 0% D_2_O to around 7 hours at 100% D_2_O (Fig. 1B). The observed growth retardation was not caused by cell death, since cell viability was only marginally affected after prolonged incubation with D_2_O (Fig. 1D).

**Figure 1.**
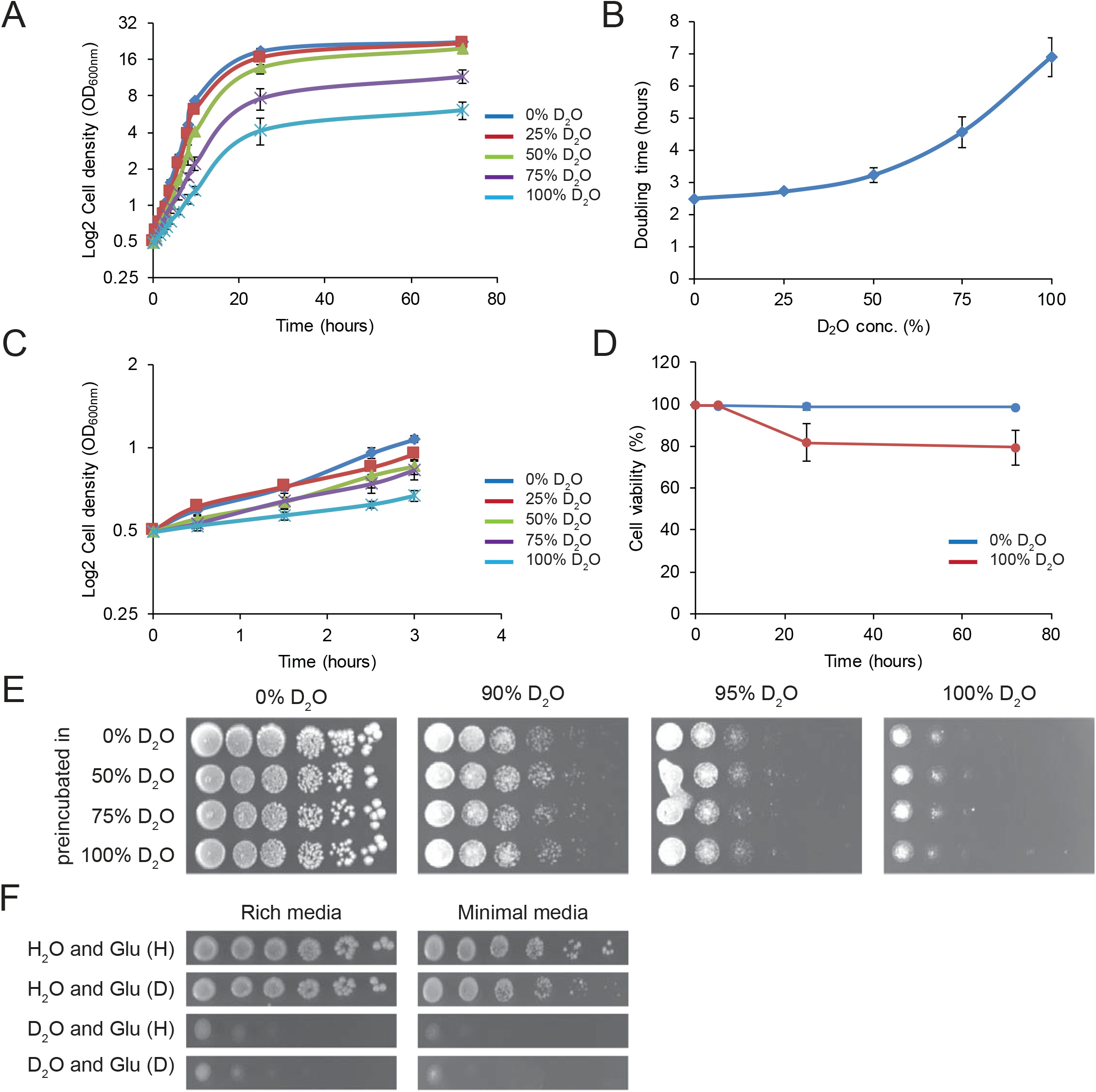
D_2_O inhibits cell growth. (A) Cell growth of wild type (no marker) *S. pombe* cells in rich medium was followed by measuring the turbidity of the culture at the indicated concentrations of D_2_O. The error bars indicate the standard deviation (n=3). (B) From growth experiments as shown in (A), the doubling time of wild type *S. pombe* cells in exponential phase was determined at the indicated concentrations of D_2_O. The error bars indicate the standard deviation (n=3). (C) A close up of the growth curves shown in (A) during the first three hours of culturing at the indicated concentrations of D_2_O. Note that the growth retardation caused by D_2_O occurs rapidly. The error bars indicate the standard deviation (n=3). (D) The viability of wild type cells grown at 0% and 100% D_2_O was determined by VitaBright staining and quantified by automated fluorescence microscopy. The error bars indicate the standard deviation (n=3). (E) The growth of wild type cells, pre-incubated for 5 hours at 0, 50, 75 or 100% D_2_O in liquid YES media, was compared by serial diluting and spotting onto solid rich media agar plates with the indicated concentrations of D_2_O at 30 °C. (F) The growth of wild type cells was compared on media prepared with D_2_O or H_2_O with either hydrogenated glucose (Glu (H)) or deuterated glucose (Glu (D)) by serial diluting and spotting onto solid rich media or synthetic minimal media agar plates at 30 °C.

Since bacteria can be adapted to growth at 100% D_2_O [17, 19], we tested if pre-incubating the cells with different amounts of D_2_O would allow for increased growth rates in the presence of D_2_O on solid rich medium. However, this was not the case for fission yeast. Hence, for all tested pre-incubations no changes in growth were observed (Fig. 1E), although we noted that when grown on solid media, wild type cells eventually formed colonies even at 100% D_2_O.

The observed effects of D_2_O may be caused by either incorporating deuterium into macromolecules or by replacing the solvent (H_2_O) with D_2_O or by a combination of the above. Thus, we compared the growth of cells on media prepared with normal glucose or deuterated glucose (glucose-C-d_7_). On rich media, there was no obvious effect of exchanging glucose with deuterated glucose (Fig. 1F). Since amino acids are plentiful in this medium, significant protein deuteration of non-solvent-exchangeable protons is not expected. On minimal medium, a slight growth inhibitory effect was observed (Fig. 1F). Since glucose here also serves as a carbon source for amino acid synthesis, the slightly reduced growth is, at least in part, likely to be ascribed to deuteration of proteins and other biomolecules. However, since the effect of deuterated glucose is minor, the observed effect of D_2_O is mainly caused by the solvent replacement. Accordingly, the growth retardation observed on media prepared with both deuterated glucose and D_2_O were similar to that observed on media with hydrogenated (normal) glucose and D_2_O (Fig. 1F). We therefore conclude that the growth inhibitory effect of D_2_O largely can be attributed to the solvent isotope effect rather than an effect of deuterium-induced malfunction of various biomolecules.

### D_2_O causes gross morphological changes, increased cell wall thickness and aberrant cell septation

The reduced cell growth was accompanied by an elongated and swollen cell morphology, which was more apparent after prolonged incubation in D_2_O (Fig. 2A). Cell septation was analyzed by calcofluor staining. This revealed a clear dosage-dependent septation defect, which appeared gradually upon shifting the cells to D_2_O medium. Hence, after 5 hours in the exponential growth phase, roughly 20% of the cells in 0% D_2_O were septated, while in 100% D_2_O this was increased to 60% of the cells (Fig. 2B). After 24 hours, the cells grown in 0% D_2_O had entered stationary phase (Fig. 1A) and were therefore not septated (Fig. 2AB). However, at 100% D_2_O about 60% of the cells were still septated (Fig. 2B) and about 20% displayed multiple septa (Fig. 2AB). In addition, an increased thickness of septa was visible already after 5 hours in D_2_O, which was succeeded by unevenly shaped septa after 24 hours (Fig. 2A).

**Figure 2.**
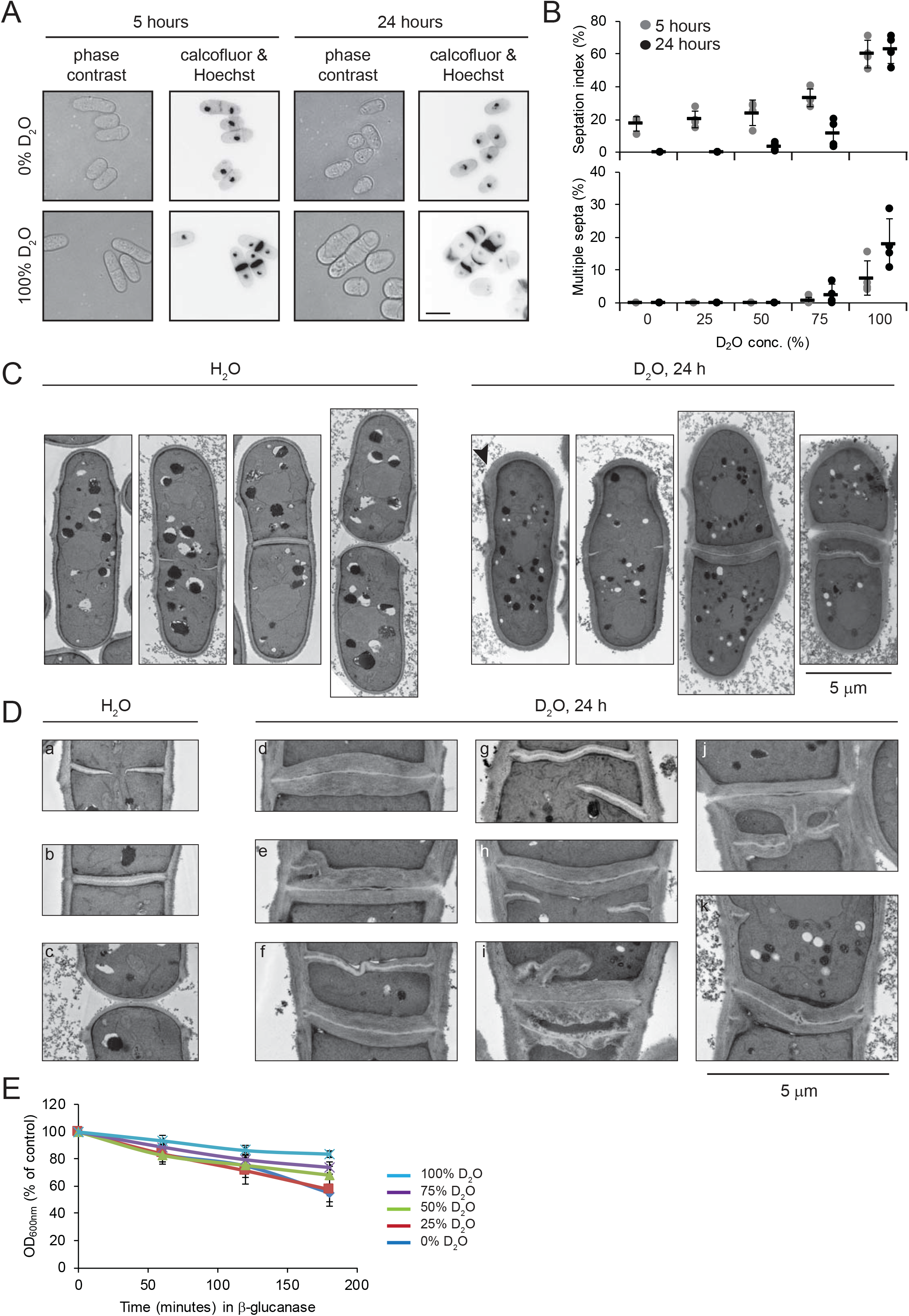
D_2_O affects cell morphology and causes cell septation defects. (A) Wild type cells grown at 0% and 100% D_2_O for the indicated times were stained with calcofluor (to mark the septa) and Hoechst (to mark the nucleus) and analyzed by microscopy. Note that the thickness of the septa is increased in the presence of D_2_O. Bar = 5 µm. (B) The percentages of septated cells (septation index) and of septated cells with multiple septa were determined after 5 and 24 hours at the indicated concentrations of D_2_O. The error bars indicate the standard deviation (n=4). (C) Transmission electron microscopy images of wild type cells after 24 hours in 0% or 100% D_2_O YES medium. Note the irregular septa and thickening of the cell wall (black arrow head). Bar = 5 µm. (D) Transmission electron microscopy images of wild type cells after 24 hours in 0% or 100% D_2_O YES medium as in panel A. Bar = 5 µm. (E) Cell lysis of wild type (no marker) cells grown at the indicated D_2_O concentrations for 24 hours was recorded over time by measuring turbidity of cultures treated with β-glucanase (in water). The error bars indicate the standard deviation (n=4).

By transmission electron microscopy we confirmed that cells grown for 24 hours in D_2_O appeared swollen and displayed thickened cell walls (Fig. 2CD). Accordingly, cells incubated for 24 hours in D_2_O-containing medium were resistant to the cell wall degrading enzyme β-glucanase, supporting that, unlike the situation for bacteria [19], D_2_O causes cell wall thickening of yeast cells (Fig. 2E). The S. pombe septum consists of primary (outer) and secondary (inner) septa that are made up of different glucans [30]. The secondary septum will eventually form the cell wall at the tip of the daughter cell. Both the primary septa (white layer) and secondary septa (grey layers flanking the primary septum) were thicker for cells grown for 24 hours in D_2_O (Fig. 2CD). In general, septum formation appeared abnormal in D_2_O. For example, in some of the D_2_O-treated cells we observed an asymmetrical septum growth from only one side (Fig. 2D, panels e, g, and k), while for the control cells, the septa always grew uniformly inwards (Fig. 2D, panels a-c). Once the septa formed, their closure appeared normal, but for the cells in D_2_O, septum material continued to build up, leading to strongly thickened secondary septa (Fig. 2D, panels d-k). Since the primary septa were stabilized in D_2_O, the cells failed to detach and remained associated with sister cells from previous generations (Fig. 2D, panel j). However, rather than reinitiating another cell cycle and forming a chain of cells, most of the multiseptated cells appeared to have formed the second or third septum near the original septum (Fig. 2D, panels e, f, and j), and not between nuclei, indicating that no further nuclear division occurred. This may suggest that D_2_O affects the septation initiation network [31]. Moreover, in some cells we observed that the second or third septum seemed to branch from the original septum, or to grow perpendicular to it (Fig. 2D, panels i and j).

### D_2_O affects both the actin and tubulin cytoskeleton

To determine if the cytoskeleton was affected, wild type cells carrying the LifeAct-GFP actin marker and GFP-tagged α-tubulin (Atb2) were exposed to D_2_O and analyzed by fluorescence microscopy. In *S. pombe* the actin cytoskeleton typically appears as cortical patches located mainly at the growing cell tips during interphase and as a contractile ring, which forms during mitosis [32]. The tubulin cytoskeleton is composed of cables that extend between the cell ends in interphase and as a dense spindle between the sister chromatids during mitosis [32].

Upon treatment with D_2_O, we observed that the actin patches were no longer concentrated at the cell tips, but rather appeared more evenly distributed throughout the cells (supporting information, Fig. S1A). The formation of the contractile ring, however, appeared normal (supporting information, Fig. S1A). In response to D_2_O, the tubulin cables were shorter, more numerous, and in some cells perpendicular rather than parallel to the cell (supporting information, Fig. S1B). However, similar to the situation with the actin cytoskeleton in mitosis, the mitotic spindle appeared unaffected by D_2_O (supporting information, Fig. S1B). Thus, unlike the situation in mammalian cells [16], the mitotic spindle was formed and appeared normal even after prolonged incubation in D_2_O.

### D_2_O inhibits meiosis

To test whether the solvent replacement affected cell cycle progression, cells exposed to different concentrations of D_2_O were analyzed for DNA content by DNA staining and automated microscopy. In *S. pombe*, cytokinesis normally occurs in S phase and cells in G_1_ and S phase are therefore binuclear and, by DNA staining, indistinguishable from the cells in G_2_. Accordingly, the DNA content profile presents as a single 2N peak. After five hours of exponential growth in D_2_O, the DNA peak remained unchanged. However, we note the appearance of a small population with 4N DNA content (supporting information, Fig. S2A), which may represent cells that have re-replicated DNA without prior mitosis or cells that have failed to undergo cytokinesis. After 24 hours, when the cultures with low concentrations of D_2_O had entered stationary phase with some cells arrested in the G_1_ phase, the 1N peak was still not evident for cells grown at 75% or 100% D_2_O (supporting information, Fig. S2A). This may indicate a reduced progression through mitosis, whereas the sharp 2N peak indicates that the cells spend a relatively short time period in S phase, suggesting that progression through S phase is not strongly affected. To more directly test whether mitosis was affected, the number of cells displaying spindles was determined using the Atb2-GFP strain. After 24 hours in D_2_O, we observed a small increase in the number of cells with spindles (supporting information, Fig. S2B), indicating that D_2_O only causes a slightly slowed progression through mitosis, and the reduced cell growth of *S. pombe* cells in D_2_O is therefore most likely caused by an overall slowing of the cell cycle.

To test if meiosis was affected, a wild type strain (*h^90^*) was starved for nitrogen in the presence of H_2_O or D_2_O. After two days, multiple asci were evident for the cells grown on the H_2_O control medium, whereas for cells incubated on D_2_O medium we were never able to detect any asci (supporting information Fig. S3). Accordingly, iodine stained the starch in spore walls dark for the cells on the H_2_O medium, but failed to stain the cells on the D_2_O medium (supporting information Fig. S3AC), suggesting that zygote formation and/or meiosis is severely inhibited by D_2_O. Using diploid cells, we observed numerous asci when the cells were incubated in H_2_O, but only a few asci for cells on D_2_O medium (supporting information Fig. S3BC). Collectively, these data indicate that D_2_O inhibits mating and meiosis. An obvious hypothesis is that the mating defect, at least in part, is caused by the D_2_O-induced cell wall thickening.

### D_2_O blocks glucose metabolism through inhibition of glucose-6-phosphate isomerase

Since the isotope effects of D_2_O are likely to disturb metabolism [33], we decided to investigate whether D_2_O affects the kinetics of enzymes in the glycolytic pathway on a short timescale. This was accomplished using hyperpolarized NMR (signal enhanced nuclear magnetic resonance) [34] which allows detailed metabolic mapping of glycolysis. A ^13^C-labelled glucose tracer was fed to live cells in either 100% D_2_O or 100% H_2_O MES buffer for 2 or 10 minutes, after which the metabolism was quenched in acid and the soluble metabolites extracted. This extract was then hyperpolarized and analyzed with single spectrum ^13^C-NMR. Examples of an NMR spectra from cells fed with the glucose tracer in D_2_O and H_2_O, respectively, are shown in Fig. 3A. The crowded spectral region from 60-100 ppm contains signals from the substrate glucose. The singlet signal at 24 ppm originates from an added standard, which allows quantification and direct comparison between the metabolite signals in the different spectra [35]. This comparison clearly revealed that several metabolic changes appear when *S. pombe* is fed glucose in D_2_O. In particular, we noted the accumulation of a large pool of α-glucose-6-phosphate (α-G6P) (Fig. 3A, blue insert), suggesting that the α anomer is being metabolised more slowly in D_2_O. Whereas β-G6P is readily used as a substrate in the pentose phosphate pathway (PPP), α-G6P must be converted to β-G6P for use in PPP or to β-fructose-6-phosphate (β-F6P) for use in glycolysis. A single enzyme, glucose-6-phosphate isomerase (GPI, EC 5.3.1.9), is responsible for both types of isomerase activities (https://www.genome.jp/kegg/). The PPP was indeed active, as demonstrated by the signals from 6-phosphogluconate (6PGA) (Fig. 3A, red insert). The 6PGA pool was approximately twice as large in D_2_O as in H_2_O. That the activity of PGI could be negatively affected by the exchange of H_2_O for D_2_O is plausible since the mechanism of catalytic isomerization involves several proton transfer steps [36].

**Figure 3.**
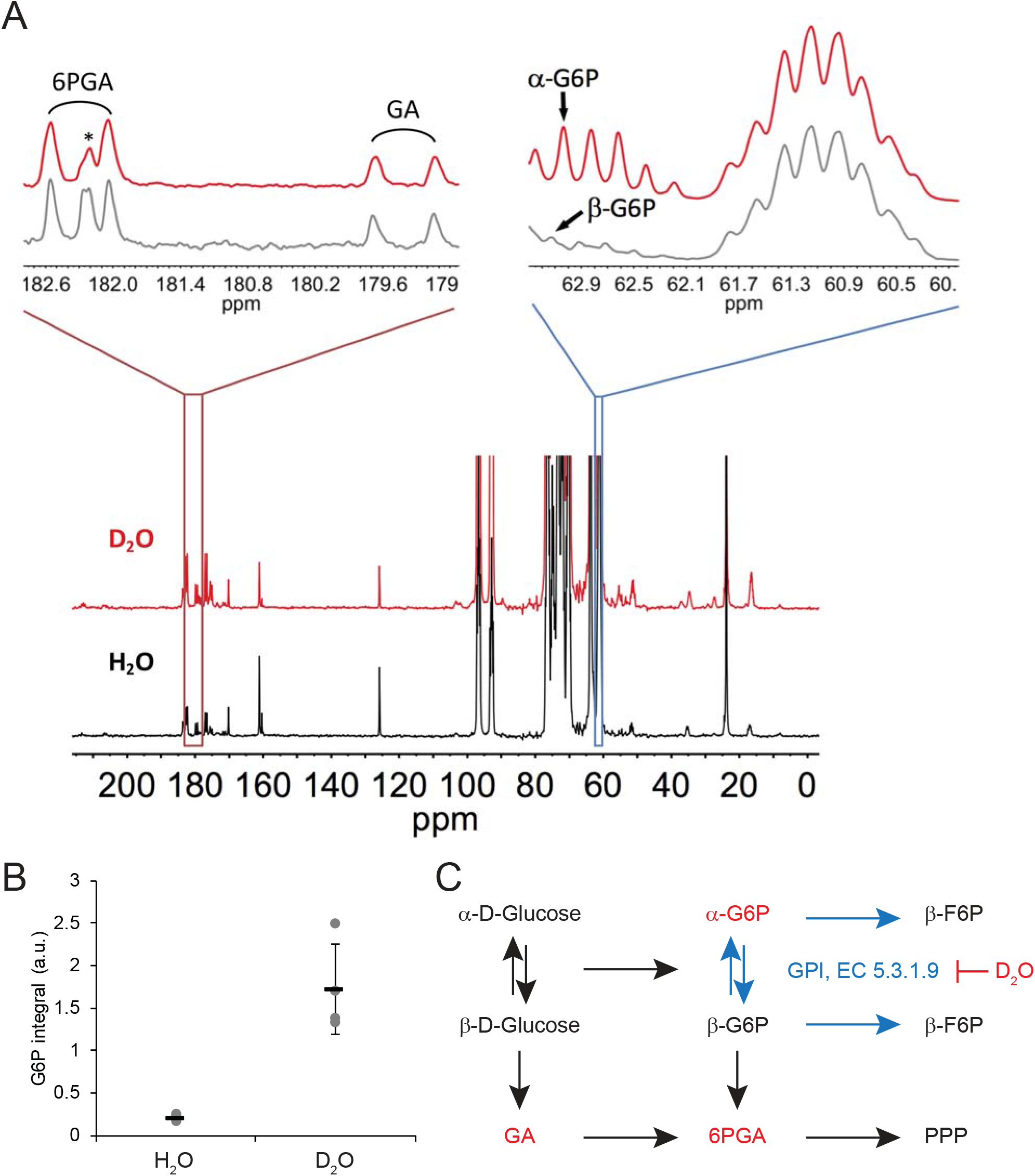
D_2_O inhibits glucose metabolism. (A) Hyperpolarized ^13^C NMR spectrum of cellular metabolite extract following incubation with the isotope labelled glucose tracer for 10 minutes in D_2_O buffer (red spectrum) and H_2_O buffer (black spectrum). Red insert shows a zoom-in on the carbonyl region showing gluconate (GA) and 6-phosphogluconate (6PGA). The asterisk marks an unidentified metabolite. Blue insert shows a zoom-in on the aliphatic sugar region, revealing a multiplet centred at 61.2 ppm originating from 6-^13^C-glucose. The multiplet at 62.8 ppm originates from the α-form of glucose-6-phosphate (α-G6P) in the spectrum from the D_2_O experiments (red) and from the β-form of glucose-6-phosphate (β-G6P) in the spectrum from the H_2_O experiments (black). (B) Quantitative difference between accumulated glucose-6-phosphate (G6P) signal in D_2_O and H_2_O exposed cells following incubation with ^13^C_6_-d_7_-labelled glucose for 10 minutes. The error bars indicate the standard deviation (n=4). (C) Glucose metabolic pathways when *S. pombe* cells are exposed to D_2_O and H_2_O. Metabolites identified in panel A are highlighted in red. Glucose in its β-form is in both D_2_O and H_2_O buffers converted into gluconate (GA) and further into 6-phosphogluconate (6PGA) in the pentose phosphate pathway (PPP). In D_2_O, the α-form of glucose-6-phosphate (α-G6P) is accumulating whereas the β-form is metabolized more rapidly. Contrary, in H_2_O, no accumulation of α-G6P can be detected. An explanation for this difference is an altered activity of glucose-6-phosphate isomerase (GPI, EC 5.3.1.9) when the cells are exposed to D_2_O. This enzyme catalyzes the isomerization reactions between α-G6P and β-G6P, α-G6P and β-F6P, and β-G6P and β-F6P (blue reactions).

The doublet signal at about 179.4 ppm, assigned to gluconate (GA) (Fig. 3A), confirms that the gluconate shunt is active in *S. pombe* cells [37, 38]. In 30-sec. time-resolved data (supporting information, Fig. S4) only gluconate could be detected and the gluconate shunt seemed to be unaffected by D_2_O.

The accumulation of α-G6P in the D_2_O-exposed cells is already apparent after 2 minutes of incubation with the glucose tracer (supporting information, Fig. S5). However, in order to facilitate quantification and comparison of the amount of G6P under the two studied conditions, exposure times of 10 minutes were used, revealing an about 8 times larger pool of G6P in D_2_O exposed cells (Fig. 3B). In summary, glycolysis as well as the pathway leading to PPP are affected negatively by an inhibition of α-G6P isomerization in *S. pombe* cells exposed to D_2_O (Fig. 3C). We conclude that, under these conditions, *S. pombe* cells use a highly active gluconate shunt to produce PPP metabolites from glucose, which allow the cells to bypass GPI.

### D_2_O induces a cellular stress response

Recently, it was reported that deuterium causes dramatic changes in the *E. coli* proteome [19]. To determine if the D_2_O-triggered growth inhibition was accompanied by differential gene expression, total RNA was purified from *S. pombe* cells under each condition in quadruplicates, and analyzed by next-generation sequencing. The four repeats of each condition clustered in separate groups (Fig. 4AB). In total, we could assign read counts to 6992 of 7015 annotated genes in the *S. pombe* genome. By setting a significance cut-off of absolute log2 fold change ≥2 between the groups and adjusted *p* values ≤ 5% (false discovery rate, FDR), we identified 178 genes that were significantly up-regulated, while 38 genes were significantly down-regulated after five hours with D_2_O. After 24 hours in D_2_O 284 genes were significantly up-regulated and 51 genes were significantly down-regulated. The identified differentially expressed genes are listed in the supporting material (supporting material file 1). The sequencing data has been uploaded to Gene Expression Omnibus (https://www.ncbi.nlm.nih.gov/geo/; accession no. GSE119785). To further test the quality of the dataset, we quantified the mRNA by real-time PCR of selected up-regulated genes. In agreement with the RNA sequencing, the expression of these genes was up-regulated (supporting information, Fig. S6).

When analyzing the differentially expressed genes, we noted a strong D_2_O-dependent down-regulation of several cytosolic and mitochondrial tRNAs (Table S1). This is in line with earlier observations in *E. coli* [39] and the reduced abundance of translational proteins recently observed by proteomics [19]. Comparison of our datasets with previous transcriptomic analyses in fission yeast (Chen et al., 2003; Poulsen et al., 2017; Vjestica et al., 2013) indicated that D_2_O causes a general stress response in the cells. Thus, the D_2_O-triggered gene expression pattern resembled those observed upon heat shock, deletion of the Hsf1 transcriptional repressor Mas5 or the Hsp70 chaperone Ssa2, or overexpression (OE) of the co-chaperone Bag101 (Fig. 4CD). We therefore tested if prior activation of the stress response pathway conferred any fitness advantage to the cells in the presence of D_2_O. To this end, wild type cells were given a 30 minute heat shock at 43 °C immediately prior to testing growth in the presence of D_2_O on solid media. However, rather than protecting the cells from D_2_O, this revealed a slight, but reproducible, decreased D_2_O tolerance (Fig. 4E), suggesting that induction of molecular chaperones, oxidoreductases and other stress-relieving proteins are unable to counteract the adverse effects of D_2_O. Similarly, applying oxidative stress via a pre-treatment of cells for 1 hour with 0.7 mM H_2_O_2_ also did not provide protection from D_2_O (Fig. 4F). Conversely, pre-incubating cells for 5 hours with 100% D_2_O led to an increased survival of cells after a severe heat shock (Fig. 4G) and an increased growth in the presence of H_2_O_2_ (Fig. 4H). This suggests that the D_2_O-triggered induction of the stress-response genes leads to increased tolerance towards heat-shock and oxidative stress conditions.

**Figure 4.**
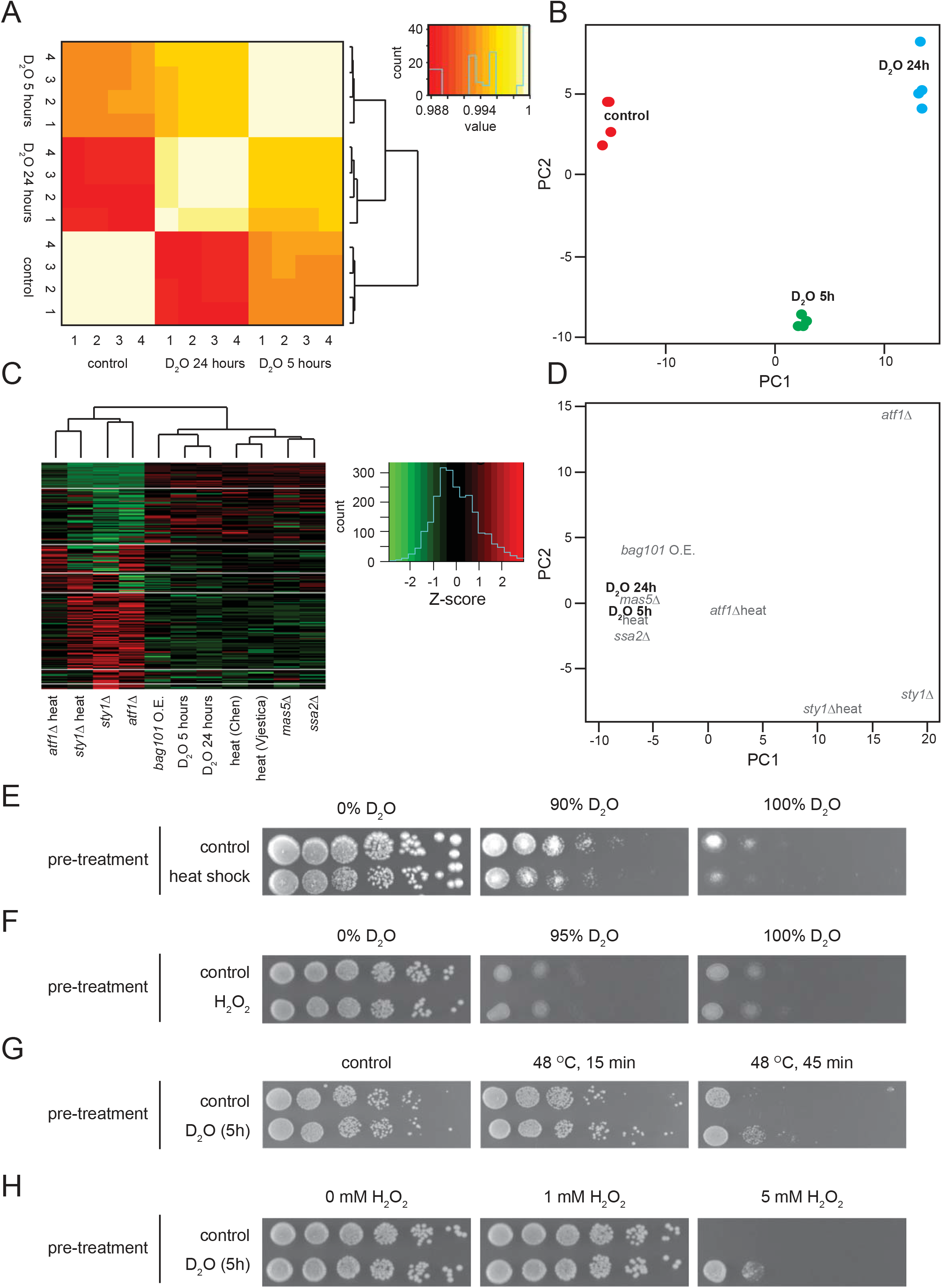
RNA sequencing of cells treated with D_2_O. (A) Total RNA was sequenced from wild type (no marker) cells that were either untreated (0% D_2_O) or treated with 100% D_2_O for 5 or 24 hours in quadruplicates. Heatmap showing hierarchical clustering on the Euclidian distances between full set of gene counts for all samples (after DESeq2’s own log transformation and variance stabilization). (B) Plot showing the first two principal components from a principal component analysis (PCA) on the full set of gene counts. (C) Heatmap of CESR genes as defined by Chen *et al*. for D_2_O regulated genes, *mas5*Δ, *ssa2*Δ, heat shock [97], *sty1*Δ, *atf1*Δ [60], and *bag101* overexpression (OE) [98]. The combined datasets were quantile normalized to make them comparable. Color key including value frequency is shown next to the heatmap. Note that the D_2_O response resembles the heat stress, *mas5*Δ, *ssa2*Δ and *bag101* OE response, but not that in the other strains. (D) Plot showing the first two principal components from a principal component analysis (PCA) on the same data shown in (C). (E) The growth at 30 °C of wild type (no marker) cells that were either untreated (control) or subjected to a 30 minute heat shock at 43 °C was compared in the presence of the indicated concentrations of D_2_O by serial diluting and spotting onto solid rich media. (F) The growth at 30 °C of wild type cells that were either untreated (control) or exposed to 0.07 mM H_2_O_2_ for 1 hour, was compared in the presence of the indicated concentrations of D_2_O by serial diluting and spotting onto solid rich media. (G) Wild type cells that were either untreated (control) or subjected to 5 hours incubation in rich medium containing 100% D_2_O were challenged with the indicated heat shock conditions prior to spotting onto solid rich media at 30 °C. (H) The growth of wild type cells that were either untreated (control) or subjected to 5 hours incubation in rich medium containing 100% D_2_O was compared in the presence of the indicated concentrations of H_2_O_2_ by serial diluting and spotting onto solid rich media at 30 °C.

**Table 1.** Gene deletions scored as D_2_O hypersensitive in high-throughput screen

| Systematic ID | Gene name | Function |
| --- | --- | --- |
| SPAC13G7.03 | *upf3* | up-frameshift suppressor 3 family protein |
| SPBC1539.08 | *arf6* | ADP-ribosylation factor, Arf family Arf6 |
| SPAC2F3.02 | - | ER protein translocation subcomplex subunit |
| SPAC328.01c | *msn5* | karyopherin/importin beta family nuclear import/export signal receptor |
| SPAPB2B4.02 | *grx5* | mitochondrial [2Fe-2S] cluster assembly & transfer glutaredoxin Grx5 |
| SPAC4D7.03 | *pop2* | F-box/WD repeat protein Pop2 |
| SPCC970.10c | *brl2* | ubiquitin-protein ligase E3 Brl2 |
| SPBC18H10.06c | *swd2* | Set1C complex subunit |
| SPBC19C7.02 | *ubr1* | N-end-recognizing protein E3 Ubr1 |
| SPAC17H9.10c | *ddb1* | Cul4-RING E3 adaptor Ddb1 |
| SPAC26H5.05 | *mga2* | IPT/TIG ankyrin repeat transcription regulator of fatty acid synthesis |
| SPBC106.10 | *pka1* | cAMP-dependent protein kinase catalytic subunit Pka1 |
| SPAC6F6.01 | *cch1* | plasma membrane calcium ion import channel Cch1 |
| SPAC3H8.02 | *csr102* | Sec14 family, phospholipid-intermembrane transfer protein Csr102 |
| SPBC13G1.08c | *ash2* | Ash2-trithorax family protein |
| SPCP1E11.06 | *apl4* | AP-1 adaptor complex gamma subunit Apl4 |
| SPAC10F6.13c | *caa1* | cytoplasmic aspartate aminotransferase Caa1 |
| SPBC23G7.08c | *rga7* | RhoGAP, GTPase activating protein Rga7 |
| SPBC27B12.08 | *sip1* | Pof6 interacting protein Sip1, AP-1 accessory protein |
| SPBC21C3.20c | *git1* | C2 domain protein Git1 |
| SPAC806.07 | *ndk1* | nucleoside diphosphate kinase Ndk1 |
| SPAC644.06c | *cdr1* | NIM1 family serine/threonine protein kinase Cdr1/Nim1 |
| SPBC16E9.13 | *ksp1* | serine/threonine protein kinase Ksp1 |
| SPCC1739.14 | *npp106* | nucleoporin Npp106 |
| SPAC7D4.06c | *alg3* | dolichol-P-Man-dependent alpha(1-3) mannosyltransferase Alg3 |
| SPBC3B9.11c | *ctf1* | mRNA cleavage and polyadenylation specificity factor subunit Ctf1 |
| SPAC2F7.07c | *cph2* | Clr6 histone deacetylase associated PHD protein Cph2 |
| SPBC16E9.08 | *mcp4* | prospore membrane protein Mcp4/Mug101 |
| SPAC926.03 | *rlc1* | myosin II regulatory light chain Rlc1 |
| SPBC13G1.03c | *pex14* | peroxisomal docking protein Pex14 |
| SPBC776.15c | *kdg2* | dihydrolipoamide S-succinyltransferase Kdg2 |
| SPAC513.03 | *mfm2* | M-factor precursor Mfm2 |
| SPAC26F1.04c | *etr1* * | enoyl-[acyl-carrier protein] reductase |
| SPBP8B7.11 | *nxt3* | ubiquitin protease cofactor Nxt3 |
| SPAC12B10.12c | *rhp41* | DNA repair protein Rhp41 |
| SPCC777.10c | *ubc12* | NEDD8-conjugating enzyme Ubc12 |
| SPAC4H3.13 | *pcc1* | EKC/KEOPS complex subunit Pcc1 |
| SPAC5D6.05 | *med18* | mediator complex subunit Med18 |
| SPCC970.06 | *erv29* | COP II adaptor Erv29 |

\*: Upregulated in RNA seq. at 24 hours in D<sub>2</sub>O.

### Screening the S. pombe gene deletion library for D_2_O sensitive or D_2_O resistant mutants

Previous studies in budding yeast have shown that heavy water sensitivity is a conditional phenotype and that a loss-of-function mutant in ASP5, encoding an aspartate aminotransferase, is hypersensitive to D_2_O [20]. As a next step, we therefore explored the genetic requirements for D_2_O tolerance by individually screening 3233 different mutants from the fission yeast haploid gene deletion collection for hypersensitivity to D_2_O on solid media.

Stationary phase cells were replica plated in 384-pin format onto YES plates with 0% or 100% D_2_O. This concentration of D_2_O was used because it completely blocked growth of a deletion strain in the ASP5 orthologue in fission yeast, *caa1*, but also impaired the growth of wild type cells (Fig. 5A), so both D_2_O hypersensitive and resistant mutants could be scored. We selected mutants with reduced growth on D_2_O medium, but excluded mutants with slow growth on 0% D_2_O medium. This led to 39 mutants scored as D_2_O sensitive (Table 1). We noted that the positive control, *caa1*Δ, was among these. We also isolated one mutant, *pmk1*Δ, deleted for the gene encoding the Pmk1 MAP kinase that appeared resistant to D_2_O.

**Figure 5.**
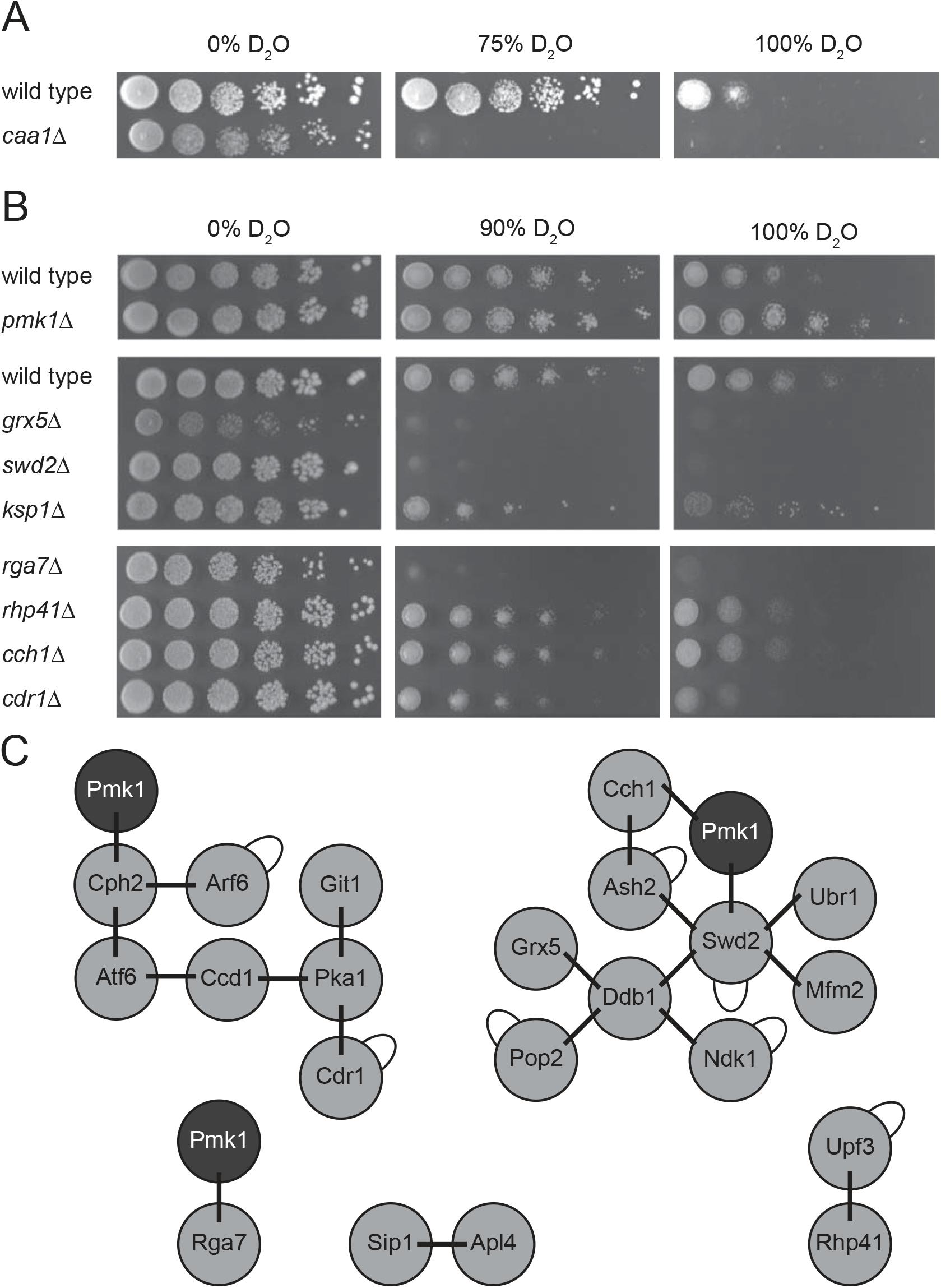
D_2_O sensitive and D_2_O resistant S. pombe mutants. (A) The growth of a 3 marker wild type strain and the otherwise isogenic *caa1*Δ were compared in the presence of the indicated concentrations of D_2_O by serial diluting and spotting onto solid rich media at 30 °C. (B) The growth of a 3 marker wild type strain and the otherwise isogenic deletion mutants were compared in the presence of the indicated concentrations of D_2_O by serial diluting and spotting onto solid rich media at 30 °C. (C) Schematic representation of pairwise genetic or physical interactions listed in the BioGRID database (version 3.4.164) between the deleted genes/proteins in the isolated D_2_O hypersensitive strains (grey) and D_2_O tolerant strain (Pmk1, black).

To assess the validity of the screen, we tested selected D_2_O sensitive mutants and the *pmk1*Δ strain in growth assays on solid media. This revealed that the *pmk1*Δ strain was slightly resistant to D_2_O, while the selected sensitive mutants all displayed a reduced D_2_O tolerance (Fig. 5B). In addition, using the BioGRID database (https://thebiogrid.org/) [40] we noted several previously identified genetic interactions between many of the D_2_O hypersensitive/resistant strains, including ones between the D_2_O tolerant *pmk1*Δ strain and the D_2_O sensitive *cch1*Δ [41, 42], *rga7*Δ [43, 44], *cph2*Δ [45] and *swd2*Δ [45] (Fig. 5C).

### D_2_O activates the cell wall integrity pathway

To obtain further insight into the biological effects of the solvent replacement, we set up a screen for spontaneous D_2_O resistant mutants. Wild type cells were inoculated in 100% D_2_O rich medium in 20 separate tubes and incubated at 30 °C. After about one week, slight growth was apparent in four of the tubes, suggesting that independent spontaneous mutations that allowed for D_2_O tolerance had occurred in these cultures. To avoid selecting clones, cells from each culture were spread on 100%

D_2_O plates and only one well-isolated large colony from each plate was selected. This led to the isolation of three independent heavy water resistant (hwr) mutants (*hwr1-3*), that all displayed a markedly increased growth in the presence of D_2_O, but otherwise appeared normal (Fig. 6A). The increased tolerance to D_2_O was also evident on 100% D_2_O minimal media (supporting information, Fig. S7). However, the *hwr* mutants were not resistant to the slight growth inhibitory effect conferred by deuterated glucose on minimal medium (supporting information, Fig. S7), indicating that the isolated *hwr* mutants were specifically tolerant to the solvent replacement effect.

**Figure 6.**
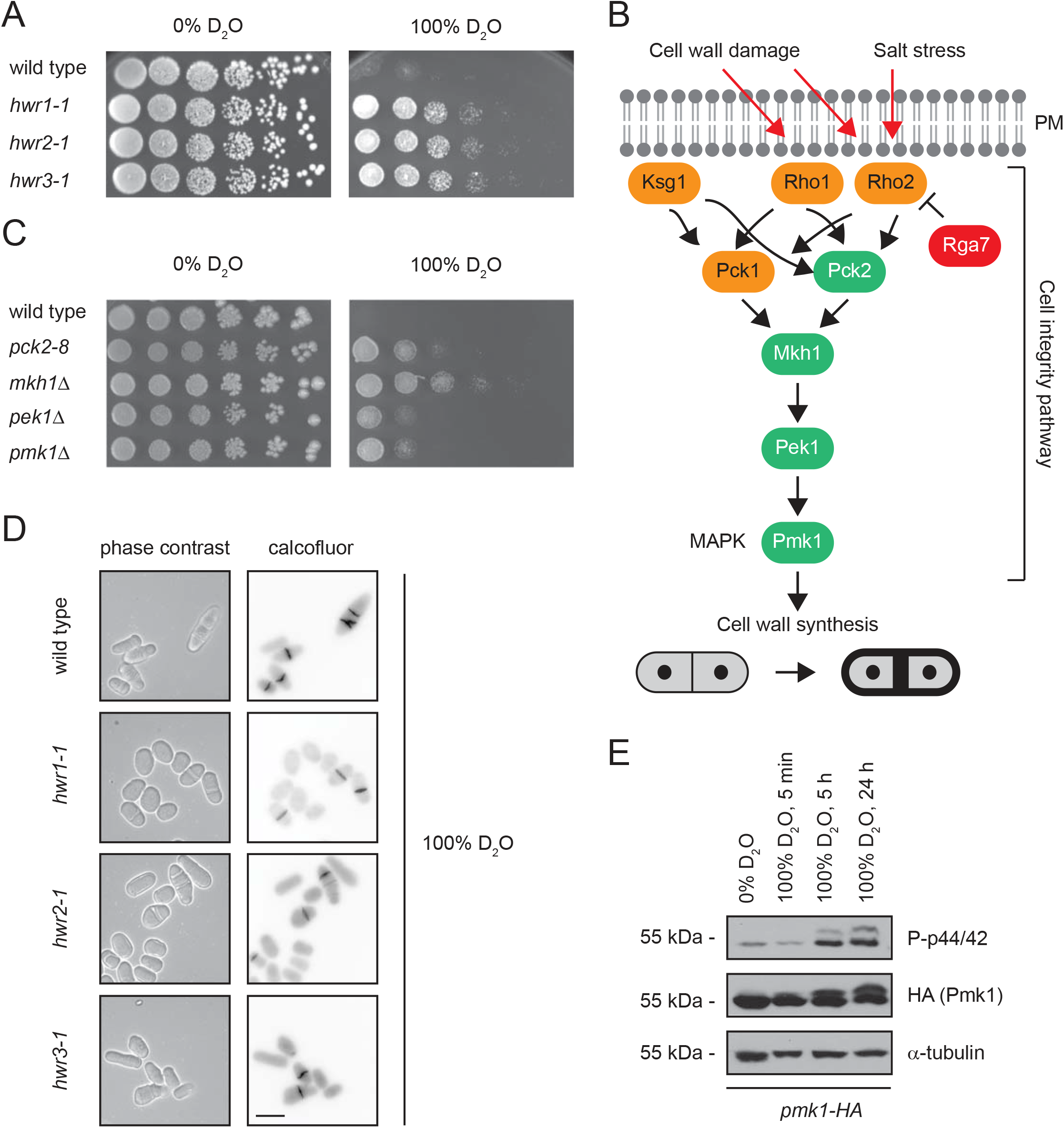
D_2_O activates the cell integrity pathway. (A) The growth of 3 marker wild type cells and the isolated *hwr* mutants was compared by serial diluting and spotting onto solid rich media agar plates. (B) The cell integrity pathway in *S. pombe* cells as described by Madrid *et al*. [47]. The pathway is known to be activated by salt stress or cell wall damaging agents (red arrows), and consists of small GTPases that activate Pck1 and Pck2. In turn, this leads to activation of Mkh1, Pek1 and finally of the MAPK Pmk1 which ultimately triggers increased cell wall synthesis. Rga7 functions as a Rho2 GAP that limits signaling [43]. The signaling components identified with mutations in the *hwr* strains are shown in green. The D_2_O hypersensitive phenotype observed for the *rga7*Δ strain is shown in red. (C) The growth of 3 marker wild type cells and the indicated mutants in the cell integrity pathway was compared by serial diluting and spotting onto solid rich media agar plates at 25 °C. (D) Wild type cells (3 marker) at 100% D_2_O for 24 hours were stained with calcofluor and analyzed by microscopy. Bar = 5 µm. (E) Whole cell lysates of wild type cells expressing HA-tagged Pmk1 were analyzed by SDS-PAGE and Western blotting using antibodies to phosphorylated Pmk1 (P-p44/42), the HA-tag on Pmk1 and, as a loading control, to α-tubulin. Note that the level of phosphorylated Pmk1 is increased in response to D_2_O.

Since the isolated *hwr* strains were the result of spontaneous mutations, as opposed to mutagenesis induced by chemicals or irradiation, we reasoned that their genome sequences were likely highly similar with the exception of the mutations causing D_2_O resistance. We therefore analyzed the *hwr* strains and their wild type parent by whole genome sequencing using Illumina HiSeq2000 high-throughput sequencing technology. The average sequence coverage was greater than 1000 reads/bp for all samples. The full sequencing datasets are deposited at the Sequence Read Archive (https://www.ncbi.nlm.nih.gov/sra; accession no. SRP161455). Comparison of obtained sequences from the *hwr* strains and the wild type control revealed a match, except for the listed changes (Table 2, and supporting material file 2).

**Table 2.** Mutect2 predicted genome sequence differences in the hwr strains

| Mutant | Chr.* | Nucleotide change* | Location | Gene | Type |
| --- | --- | --- | --- | --- | --- |
| *hwr1-1* | II | A4313777del | exon | *pek1* | frameshift |
| *hwr1-1* | III | G2337547A | exon | *trm72* | missense (G253E) |
| *hwr2-1* | I | 613724CTATGTTCAAGins | exon | *mkh1* | frameshift |
| *hwr2-1* | I | 1670566Tins | splicing | *gta2* | - |
| *hwr2-1* | I | T5448804del | 5' UTR | *SPAC1039.02#* | - |
| *hwr3-1* | I | G3273325C | ncRNA | *SPNCRNA.899* | - |
| *hwr3-1* | II | C2310230T | exon | *pck2* | missense (G870S) |

\*: Chr.: chromosome, del: deletion, ins: insertion; #: Downregulated in RNA seq. after 5 hours in D<sub>2</sub>O.

When studying the mutated genes in the *hwr1-3* strains (Table 2), we noticed that several have been linked to tRNA metabolism (*gta2* and *trm72*) and cell wall formation (*pck2*, *mkh1* and *pek1*). The kinases Pck2, Mkh1 and Pek1 were mutated in the *hwr3, hwr2* and *hwr1* strains, respectively. These kinases function as upstream activators of Pmk1 in the so-called cell integrity pathway [46, 47], which is activated by cell wall damage to upregulate synthesis of various cell wall components [48], while the Rho2 GAP, Rga7, restricts signaling through this pathway (Fig. 6B) [43, 44]. None of these genes are essential, but except for the D_2_O-sensitive *rga7*Δ and D_2_O-tolerant *pmk1*Δ, the deletion mutants were not included in the screened knock-out library. The *pek1* and *mkh1* mutations both result in frame shifts in regions encoding the respective kinase domains of Pek1 and Mkh1 that we predict will cause loss of function, while *pck2* contained a missense mutation (supporting information, Table S2). These changes were confirmed by PCR and Sanger sequencing (supporting information, Fig. S8-S10). We therefore continued to test if the D_2_O tolerant phenotype was also evident in the independent *pck2-8*, *mkh1*Δ, and *pek1*Δ strains. Indeed, in growth assays, these strains appeared more resistant to D_2_O than wild type cells (Fig. 6C), thus phenocopying the *hwr* strains. In agreement with this, the cell septation problems observed for wild type cells in D_2_O were reduced in the *hwr* strains (Fig. 6D). The conclusions above suggest that D_2_O activates the cell integrity pathway and blocking this pathway leads to an increased fitness at high concentrations of D_2_O, presumably by limiting synthesis of glycans and/or other cell wall components. To test this prediction, we directly analyzed the activation of the cell integrity pathway in response to D_2_O in a wild type strain, expressing HA-tagged Pmk1. This revealed a marked increase in the level of phosphorylated Pmk1 after 5 and 24 hours in D_2_O-containing media (Fig. 6E), which further suggests that the solvent replacement activates the cell integrity pathway. In turn, blocking the Pck2-Mkh1-Pek1-Pmk1 signaling axis, at least partially, alleviates the D_2_O-triggered cell wall and growth defects, while activating it by deletion of *rga7* increases D_2_O sensitivity (Fig. 6B).

## Discussion

In the present study, we have systematically addressed the cellular and biochemical consequences of exchanging H_2_O for D_2_O in eukaryotic cells using *S. pombe* as a model organism. The effect of D_2_O cannot directly be compared to any other insults that a cell can experience because it entails an almost complete substitution of solvent. This possibility of solvent change for biological reactions and life as such is unique. In some ways it can be compared to changing pH, which also implies global solvent effects, but contrary to shifts in pH, which are common in nature and has well-defined biological responses, a shift to D_2_O is completely abiotic and something that we can say with absolute certainty that cellular responses have not experienced before. Thus, querying the effects of D_2_O on cellular growth and metabolism highlights subtleties of the biophysical properties of H_2_O as solvent for biological processes. *A priori* one would expect that the toxic effects of replacing H_2_O with D_2_O would be the sum of multiple smaller pleiotropic effects, and it is therefore surprising that this work demonstrates that modification of a single signaling pathway is sufficient to overcome the growth defect induced by the solvent replacement.

In agreement with early studies using human cells and other model systems [5], we observed that high concentrations of D_2_O strongly reduced cell growth. In principle, this effect can be the result of deuteration of various biomolecules or the solvent exchange effect. When D is incorporated into biomolecules, this can occur either *de novo*, where D_2_O donates D atoms during *de novo* biosynthesis, or by exchange of loosely bound hydrogens in preexisting biomolecules. Hydrogen covalently bound to carbon does not exchange, and generation of C-D bonds therefore requires *de novo* synthesis. Since we only observed minor effects of exchanging glucose with deuterated glucose on minimal medium, metabolic incorporation of D into C-D bonds does not appear to strongly affect the growth of *S. pombe*. The resulting changes in stability and/or dynamics of proteins and other biomolecules, carrying C-D bonds, therefore appears negligible next to the observed solvent effects of exchanging H_2_O for D_2_O. However, upon substituting H_2_O with D_2_O, loosely-bound hydrogen atoms such as those bound to amide/amino nitrogen or hydroxy oxygen will, depending on solvent exposure, exchange with deuterium atoms in the D_2_O solvent. Here, the properties of hydrogen bonds compared to the stronger and slightly longer deuterium bonds likely become highly relevant. Hence, deuteron transfer reactions are expected to be less efficient than the corresponding proton transfer reactions. Also, the more ordered nature of D_2_O compared to H_2_O suggests that proteins and other biomolecules are less dynamic and more tightly wrapped in D_2_O than in H_2_O [49]. Accordingly, D_2_O is known to both directly (as a solvent) and indirectly (through deuteration) affect the structure and stability of proteins and other macromolecules *in vitro* [25, 50–54].

One of the direct effects of the solvent exchange that we observed was on the central metabolism, where we directly and on a very short time-scale (2 minutes) noted a clear accumulation of α-G6P, suggesting that G6P isomerase is inhibited. The solvent exchange also led to an activation of a general stress response pathway. Typically, this pathway is provoked by protein misfolding events, suggesting that solvent exchange may destabilize proteins *in vivo*. However, inducing molecular chaperones through a heat shock prior to incubation with D_2_O did not protect cells from the D_2_O-induced growth retardation. This indicates that activation of the stress response pathway is not tuned to counter the adverse effects of the solvent exchange, although it has been reported that overexpression of Hsp70 may protect budding yeast cells from D_2_O [55]. In line with chaperones being unable to protect fission yeast cells against D_2_O, we were unable to adapt *S. pombe* to growth at higher D_2_O concentrations and we did not identify any chaperone mutants as hypersensitive or resistant to D_2_O. Conversely, it was not surprising to find that pre-treating cells with D_2_O led to an increased tolerance towards heat and oxidizing conditions, which is most likely caused by the D_2_O-induced activation of the stress response pathway. The activated stress response and slow growth phenotypes are probably also connected with our observation that exposure to D_2_O caused a strong decrease in the expression of several tRNAs. In budding yeast and in bacteria it has been shown that tRNAs are degraded in response to stress conditions [56, 57], and we speculate that perhaps a similar mechanism operates in *S. pombe*.

The multiple effects, we observed for cells in the presence of D_2_O, not surprisingly, suggest that the effects of the solvent exchange are pleiotropic. Accordingly, it is easy to imagine a number of genetic defects that may cause cells to become hypersensitive to D_2_O. However, it is harder to reconcile D_2_O resistant mutants with the observed pleiotropic phenotypes, and although heavy metals, oxidizing agents and increased temperatures also induce multiple cellular effects [58–61], these conditions occur naturally and cells have therefore evolved specific transcriptional programs driving the expression of proteins that fend against such insults. Since the natural abundance of D_2_O is miniscule (<0.02% in water), any D_2_O resistant mutant must therefore be a consequence of some cellular pathway, which indirectly improves fitness in response to D_2_O. We identified a null mutant in *pmk1* (orthologue of human MAPK7) which displays increased tolerance to D_2_O. This is in line with Pmk1 acting as a MAP kinase that regulates cell wall biogenesis [44, 62, 63] and the observed D_2_O-induced cell wall and cell septation defects. Accordingly, the cell integrity pathway is activated in response to D_2_O, and when we selected for spontaneous D_2_O-tolerant strains, we recovered mutants in other components of the pathway. Moreover, we found that deletion of the Rho2 GAP, Rga7, which normally down-regulates the cell integrity pathway [43, 44], leads to a D_2_O hypersensitive phenotype. However, besides *rga7*Δ, other mutants that we scored as D_2_O sensitive have also been reported in BioGRID to display genetic interactions with components of the cell integrity pathway (supporting material file 3). Intriguingly, the D_2_O-dependent cell wall effects, we report here, resemble those observed upon deletion of the kinase Kin1, which regulates cell polarity [64, 65]. Similar to D_2_O treated cells, the *kin1*Δ strain displays positive genetic interactions with *pck2*Δ and *pmk1*Δ mutants [64].

Activation of the cell integrity pathway is in agreement with its role in mediating an appropriate response to allow survival during sudden changes in water activity [66] and earlier studies suggesting that D_2_O causes osmotic stress [67, 68]. It does, however, pose the question of how the solvent exchange, or what property of D_2_O, causes activation of the pathway? Several upstream membrane-spanning sensors of the cell integrity pathway, including Wsc1 and Mid2, have been identified in *Saccharomyces cerevisiae*. Recent studies suggest that these sensors physically couple the cell wall with the plasma membrane, and in a spring-like manner function to sense mechanical perturbations or elasticity changes in the cell wall and/or plasma membrane [69]. Accordingly, D_2_O is known to affect phospholipid membranes [70] and ion channels [68], and we speculate that this or related effects on the cell wall glycans may cause a direct activation of the cell integrity pathway.

Identifying D_2_O-resistant strains has important practical applications, since the phenotypes imposed by D_2_O severely hinder the production of deuterated proteins for structural studies [71], but also potentially for biological production of deuterated pharmaceuticals, which in some cases are superior to their protonated counterparts [72], and are therefore increasingly being considered as drug candidates [73–75]. Although bacteria can be adapted to grow in 100% D_2_O [17], growth is often poor and the bacteria quickly lose the D_2_O tolerance when returned to H_2_O media. The results presented here provide an important first step towards an understanding of how D_2_O affects cell biology and for generating D_2_O tolerant strains for production of deuterated biomolecules.

## Materials and Methods

### Heavy water and deuterated glucose

The used D_2_O (deuterium oxide) was purchased as 99.8% D_2_O/0.2% H_2_O (Sigma). Deuterated glucose (D-glucose-1,2,3,4,5,6,6-d_7_) was purchased as 97 atom % D (Sigma).

### Yeast strains and techniques

The fission yeast strains that were used in this study are all derivatives of the wild type *972h^-^* and *975h^+^* heterothallic strains that either carry no auxotrophic markers (no marker wild type), or the three auxotrophic markers *ura4-D18*, *leu1-32*, and *ade6-216* (3 marker wild type). The gene deletion strains were from the fission yeast haploid deletion library purchased from Bioneer [76]. The *LifeAct-GFP* and *atb2-GFP* strains have been described before [77, 78]. The *pck2-8* strain was kindly provided by Dr. Jeremy Hyams. The *pmk1-HA* strain was kindly provided by Dr. Marisa Madrid [47]. Growth assays in liquid YES (yeast extract with supplements) media (5 g/L yeast extract, 30 g/L glucose, 225 mg/L adenine, 225 mg/L uracil, 225 mg/L leucine) were performed at 30 °C with vigorous shaking by following the optical density (OD) at 600 nm. Growth assays on solid media, either YES media (YES medium with 20 g/L agar) or EMM2 (Edinburgh minimal media) [79, 79] were performed at 25 or 30 °C essentially as described [80]. Sporulation assays were performed using malt extract agar plates (30 g/L malt extract, 20 g/L agar). For iodine staining, solid iodine was heated in a closed beaker and the plates briefly exposed to the iodine vapor. The wild type diploid strain was constructed by mating *ade6-210 h^+^* cells with *ade6-216 h^-^* cells on malt extract and selecting for cells that could grow in the absence of adenine.

β-glucanase (Sigma) resistance was determined (in water) for cells grown for 25 hours in 100% D_2_O-based YES medium as described [81].

Cell viability was determined by staining cells using VitaBright-48 (ChemoMetec) and a NucleoCounter NC-3000 (ChemoMetec) as described by the manufacturer.

To monitor cell cycle progression, cultures treated or untreated with D_2_O, were stained with SYTOX Green and the cellular DNA content was quantified using a NucleoCounter NC-3000 (ChemoMetec) as described by the manufacturer.

### Cell imaging and microscopy

Calcofluor white (Sigma) staining was used to monitor cell septation as described previously [82]. Hoechst (Sigma) staining was used to mark the nucleus. Cells were observed on 2 % agarose pads made in EMM with H_2_O or D_2_O. Samples were examined using a Zeiss Z1 AxioObserver inverted fluorescence microscope, equipped with a motorized stage, in a temperature-controlled incubation chamber. A 100x, 1.4 NA oil immersion lens and a Cool-Snap HQ2 CCD camera (Photometrics) controlled by Axiovision software (Zeiss) was used for image capture. All image analysis was performed using ImageJ [83]. Electron microscopy was performed essentially as described [84].

### Hyperpolarized extract ^13^C-NMR

A previously published protocol [35] was followed for substrate incubation, metabolite extraction, and extract hyperpolarization. Briefly, 100 µL 120 mM U-^13^C, d_7_ glucose dissolved in either D_2_O- or H_2_O-based morpholinoethanesulfonate (MES) buffer (50 mM MES, 100 mM NaCl, pH 5.5) was added to the yeast cells and the respective samples were incubated for 2 minutes or 10 minutes at 30°C after which the metabolism was stopped by addition of 400 µL perchloric acid and the soluble metabolites were extracted according to the protocol. The freeze-dried extracts were re-dissolved in a hyperpolarization matrix and polarized at 3.35 T and 1.4 °K. Hyperpolarized extracts were hereafter dissolved in 4 mL D_2_O or H_2_O based MES-buffer pH 5.5 and injected into a 5 mm NMR tube and a single spectrum ^13^C-NMR was acquired on a 9.4 T Varian Inova spectrometer immediately with a 90 degree pulse.

NMR spectra were analyzed with the MNova software. Metabolites were identified by reference to the human metabolome database [85].

### RNA sequencing

Total RNA was isolated using the hot phenol method [86] from cells that were either untreated or treated with 100% D_2_O for 5 or 24 hours. Conversion to cDNA and sequencing was performed by the Beijing Genome Institute using the Illumina sequencing as single-end with final read lengths of 50 bp after demultiplexing. This produced 23.9-24.1 million reads per sample. The raw reads were aligned to the *S. pombe* genome using bwa mem (v0.7.12) with default settings giving an alignment rate of 98.4-98.7% for all 12 samples. Mapped reads were assigned to genes using GFOLD (‘gfold count’) followed by GFOLD statistical analysis (‘gfold diff’) to detect differentially expressed (DE) genes [87].

The significance cut-off for DE genes was set at FDR ≤ 5% and absolute log2 fold change ≥ 2. For figures the DESeq2 (R package) [88] normalized gene counts (‘rld’) were used. The *S. pombe* reference genome and gene annotation were downloaded from GenBank (date stamped 27-FEB-2015, Rel. 123, Last updated, Version 14). Supplementary data tables from Chen *et al*. (Chen et al., 2003) were downloaded from http://128.40.79.33/projects/stress/. The data from Vjestica *et al*. are primarily from the supplementary material (Vjestica et al., 2013).

Our annotation also includes rRNA, tRNA, ncRNA, snoRNA, and mitochondrial genes, that in many cases are not present in the older annotations used by Chen *et al*. (Chen et al., 2003) and Vjestica *et al*. (Vjestica et al., 2013). Thus, it was not possible to compare the expression levels of many, in particular non-coding, genes across the studies. The DESeq2 Bioconductor package in R (Anders et al., 2015) was used for statistical analyses of differentially expressed genes.

### Screening the S. pombe deletion library

The *S. pombe* haploid gene deletion library was first pinned to YES (0% D_2_O) agar plates in 384 format. Once large colonies had formed, cells were transferred to YES with 0% D_2_O and YES with 100% D_2_O agar plates in 384 format. After 8 days of incubation at 30 °C, the plates were photographed and cell growth was scored by manual inspection. All cell handling was performed using a ROTOR HDA pinning robot (Singer Instruments).

### Screening for spontaneous D_2_O-tolerant mutants and whole-genome sequencing

Wild type cells were inoculated in 1 mL 100% D_2_O YES medium in separate tubes and incubated at 30 °C. After about 1 week, when slight growth was apparent in some of the tubes, cells from each culture were spread on solid 100% D_2_O YES media plates and only one single large colony from each plate was selected for further analyses. Genomic DNA was purified by phenol-chloroform extraction as described [89]. The purified DNA from the three selected *hwr* mutants and the wild type parent was sequenced by the Beijing Genome Institute.

Illumina sequencing was applied using paired-end with final read lengths of 150 bp after demultiplexing and produced 78.4-87.5 million read pairs per sample, which leads to an average coverage >1000 reads per reference base.

The raw reads were trimmed using Trimmomatic (v0.32; default setting except: HEADCROP=10, SLIDINGWINDOW=4:30, MINLEN=30) [90]. The trimmed reads were aligned to the *S. pombe* genome (version ASM294v2.38) with ‘bwa mem’ (v0.7.15; default settings except: -M) [91] and the resulting alignments locally refined using Stampy (v1.0.32; default settings except: --sensitive--bamkeepgoodreads -M) [92] for higher mapping sensitivity. With Picard tools (v4.0.1.1) (http://broadinstitute.github.io/picard) the alignments were coordinate sorted (sortSam) and read duplicates annotated (MarkDuplicates). The alignment rate was >99% for all four samples.

To call variations (SNPs, insertions and deletions) between wild type cells and the three independent *hwr* mutants we used GATK LeftAlignIndels followed by GATK Mutect2 (v4.0.1.1; default settings except: -ploidy 1 -tumor <Mutant> -normal <Wild-type>) [93, 94]. The Mutect2 generated list of putative mutations were quality filtered with ‘bcftools filter’ (v1.3.1; settings: ‘AF[0]>0.75’). Mutations were also called on the alignment files using Freebayes (v1.2.0; default settings except: -- ploidy 1 --min-alternate-count 5 --min-alternate-fraction 0.4) followed by quality filtering by ‘bcftools filter’ (settings: ‘AO>200 && ODDS>100 && GL[*]<-100 && AC=1’) [95]. Mutect2 and Freebayes agree on the main findings. The mutations were annotated to genes (ENSEMBL, *S. pombe* version ASM294v2.38) using ANNOVAR (version: July 17 2017) [96]. Comparison to the reference genome revealed that our strains contained two mutations on chromosome II (A153954T and T153964A) in the *mal1* gene encoding maltase alpha-glucosidase resulting in a variant of Mal1 containing an R131R silent and F135I missense substitution.

Three mutations were selected for PCR validation, and multiple alignments in ClustalW comparing the annotated full length gene, the wild type PCR sequences and the *hwr* PCR sequences all verified the predicted mutations.

PCR sequencing was performed by Eurofins. The used primers were: *mkh1* forward: GATAATGTGTATGACAATGACGC, *mkh1* reverse: GCATGTCGTAAAGATTCGTG, *pek1* forward: TGACGTCAAAAGGTCAAGTG, *pek1* reverse: CTAATCAGACCAGACTTGACG, *pck2* forward: AAGCCCAGATGGTCATG, *pck2* reverse: GGATGAGTCATGACATCTTCTG.

### Electrophoresis and blotting

Whole cell lysates were prepared with TCA and glass beads as described [84]. Proteins were resolved by SDS-PAGE on 12% acrylamide gels and subsequently transferred to nitrocellulose membranes (0.2-μm pore size, Advantec, Toyo Roshi Kaisha Ltd.). The blots were blocked in PBS (133 mM NaCl, 2.7 mM KCl, 6.5 mM Na_2_HPO_4_, and 1.5 mM KH_2_PO_4_ (pH 7.4)) with 5% fat-free milk powder and 0.01% Tween 20. The antisera, diluted 1:1000, were as follows: anti-HA (clone 3F10, Roche), anti-P-p44/42 (Phospho-p44/42 MAPK #9101, Cell Signaling Technology), and anti-α-tubulin (clone TAT1, Abcam). Secondary HRP-conjugated antibodies were from Dako Cytomation. Blots were developed using the Pierce ECL Plus detection kit (Thermo Scientific).

## Acknowledgements

The authors thank Dr. Marisa Madrid and Dr. Jeremy Hyams for sharing yeast strains, Dr. Shiraz Shah for help with the RNA sequencing and Dr. Genevieve Thon for help with screening the deletion library. In addition, we thank Mrs. Anne-Marie Lauridsen for expert technical assistance, and Dr. Kresten Lindorff-Larsen, Dr. Olaf Nielsen, Dr. Erik Boye, Dr. Michael A. Sørensen, Dr. Sofie V. Nielsen, and Dr. Martin Willemoës for helpful discussions and comments on the manuscript.

## Author contributions

C.K., C.H., M.K., S.K.G., M.H.L., I.J. conducted the experiments. C.K., C.H., I.J., M.K., M.H.L, B.B.K, J.V.J., and R.H.P. analyzed the data. C.K., J.V.J., H.N.B., M.H.L., J.R.W. and R.H.P. designed the experiments. J.R.W. and R.H.P. conceived the study. C.K., J.R.W. and R.H.P. wrote the paper.

