## supporting information for "Mutations in a single signaling pathway allow growth on a different solvent than water"

### *Supporting material*


*Supporting figure S1 (Effects of $D_2O$ on the cytoskeleton):* *p2*

*Supporting figure S2 (The effect of $D_2O$ on mitosis):* *p4*

*Supporting figure S3 ($D_2O$ inhibits mating and meiosis):* *p5*

*Supporting figure S4 (Hyperpolarized time-resolved $^{13}C$-NMR spectra):* *p6*

*Supporting figure S5 (Single $^{13}C$ NMR spectrum of cellular metabolite extract):* *p7*

*Supporting figure S6 (Real-time PCR on selected differentially expressed genes):* *p8*

*Supporting figure S7 (Growth assays of hwr strains on deuterated glucose):* *p9*

*Supporting figure S8 (PCR sequence confirmation of hwr1-1):* *p10*

*Supporting figure S9 (PCR sequence confirmation of hwr2-1):* *p11*

*Supporting figure S10 (PCR sequence confirmation of hwr3-1):* *p12*

*Supporting table S1 (Differentially expressed tRNAs):* *p13*

*Supporting table S2 (Amino acid changes in affected kinases in the hwr strains):* *p14*

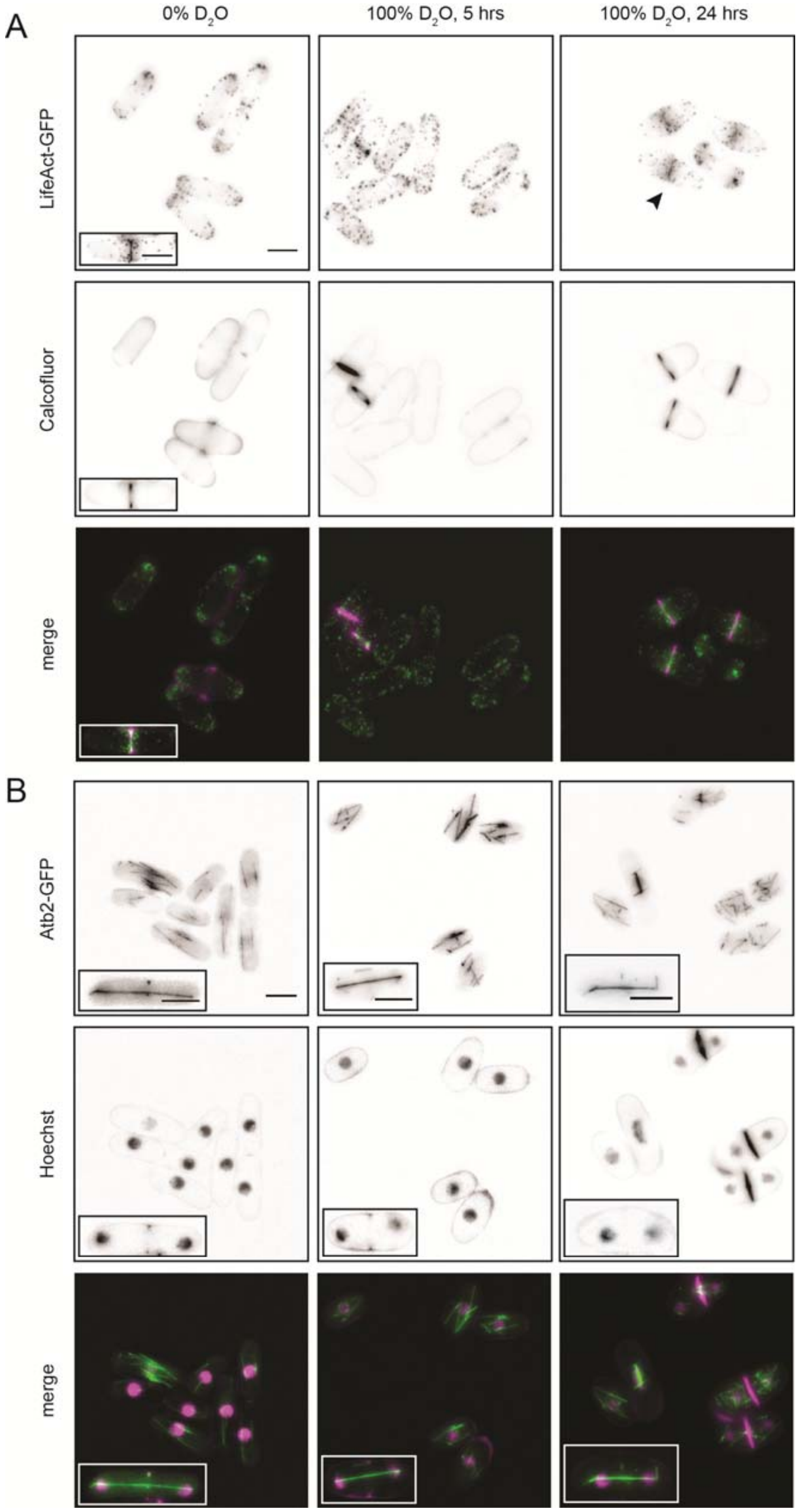
A
0% D2O
100% D2O, 5 hrs
100% D2O, 24 hrs
LifeAct-GFP
Calcofluor
merge
B
Atb2-GFP
Hoechst
merge

**Figure S1.** *Effects of $D_2O$ on the cytoskeleton.* (A) Wild type cells carrying the LifeAct-GFP actin marker cultured in 0% and 100% $D_2O$ EMM medium for the indicated times were stained with calcofluor (to mark the septa) and analyzed by microscopy. Note that the contractile ring appears normal (insert and arrow head). Bar = 5 μm. (B) Wild type cells carrying GFP-tagged α-tubulin Atb2 cultured in 0% and 100% $D_2O$ EMM medium for the indicated times were stained with Hoechst (to mark the nucleus) and analyzed by microscopy. In some cells Hoechst also stained the septa. Note that the mitotic spindle appears normal (inserts). Bar = 5 μm.

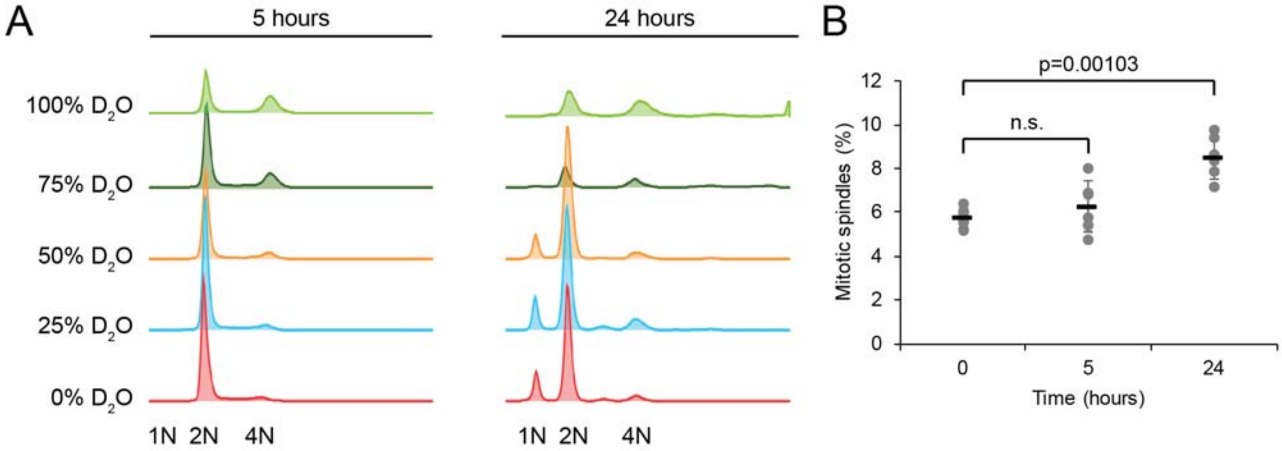


**Figure S2.** *The effect of* $D_2O$ *on mitosis.* (A) DNA content analyses. Wild type cells grown at the indicated concentrations of $D_2O$ were DNA stained and analyzed by automated fluorescence microscopy. The positions of the 1N and 2N DNA content peaks are marked. (B) The percentage of Atb2-GFP cells in displaying spindles was determined by fluorescence microscopy after 0, 5 and 24 hours in 100% $D_2O$ by counting cells. The error bars indicate the standard deviation (n=6). Student's t-test p-value is shown; n.s.. not significant.

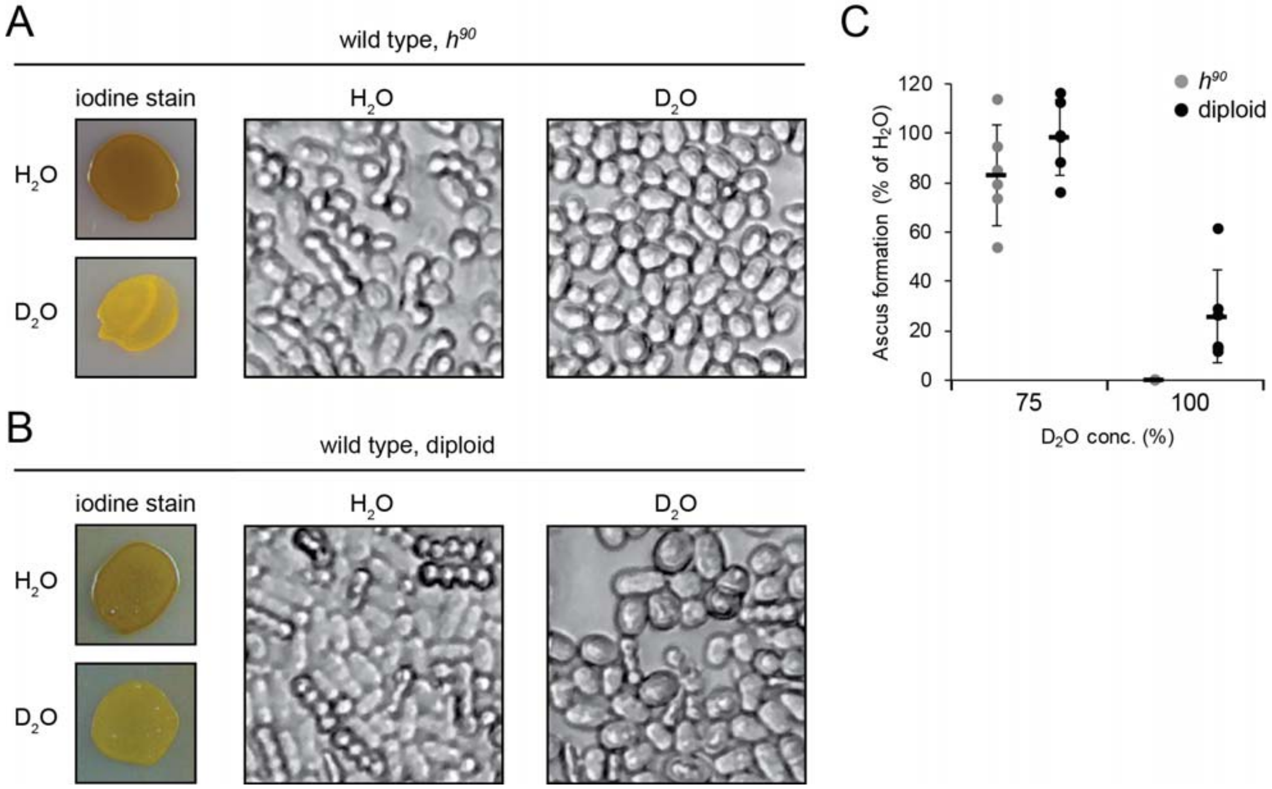


**Figure S3.** *$D_2O$ inhibits mating and meiosis.* (A) Wild type $h^{90}$ cells were plated on malt extract (to induce mating) with 0% or 100% $D_2O$, and after 48 hours analyzed by iodine staining and microscopy. Note the presence of asci containing four spores for the cells cultured in $H_2O$, and the absence of asci for cells cultured in $D_2O$. (B) Wild type diploid cells were plated on malt extract with 0% or 100% $D_2O$, and after 24 hours analyzed by iodine staining and microscopy. (C) Ascus formation was quantified by counting the number of asci. The error bars indicate the standard deviation (n=6).

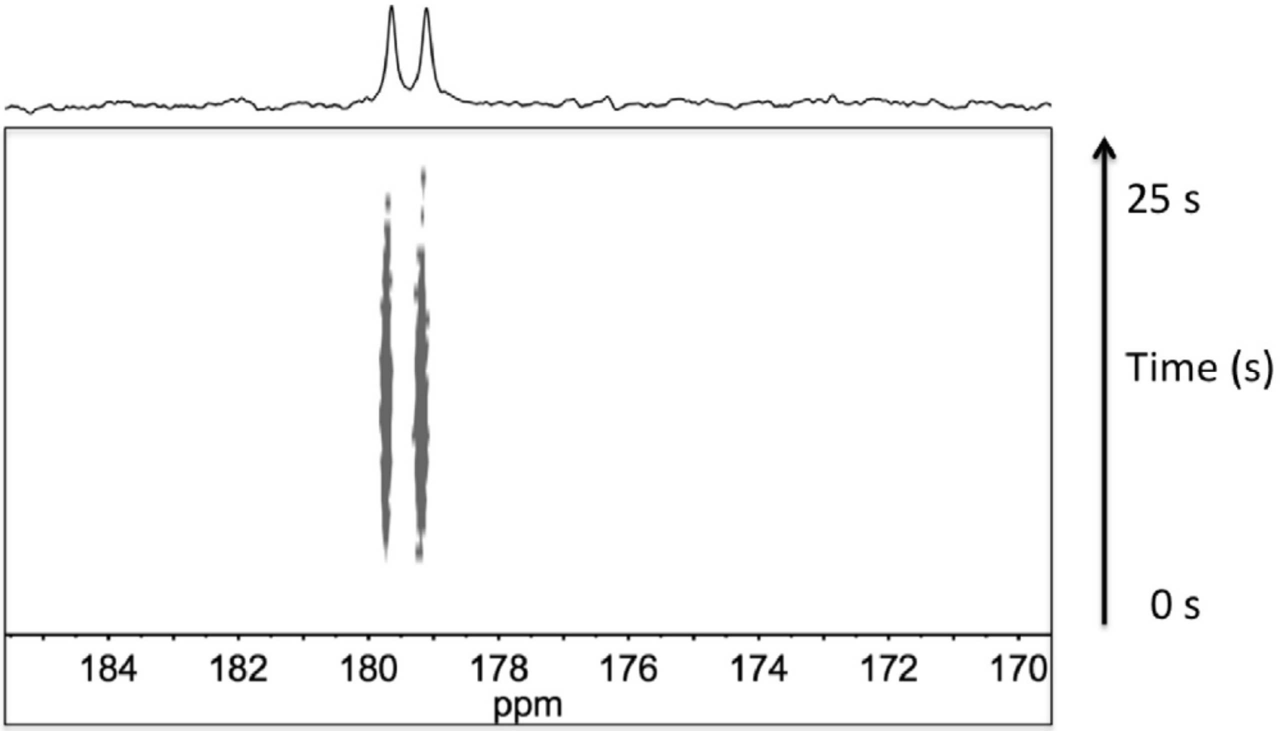


**Figure S4.** *Hyperpolarized time-resolved $^{13}C$-NMR spectra of 1-$^{13}C$-gluconate*. A hyperpolarized NMR signal is seen developing over 25 seconds at 179.4 ppm. In the top panel a sum of 50 spectra in the dynamic series is shown. Since 1-$^{13}C$-gluconate originates from uniformly $^{13}C$ labelled glucose the signal is split due to J-coupling with neighbouring $^{13}C$-enriched carbon atom.

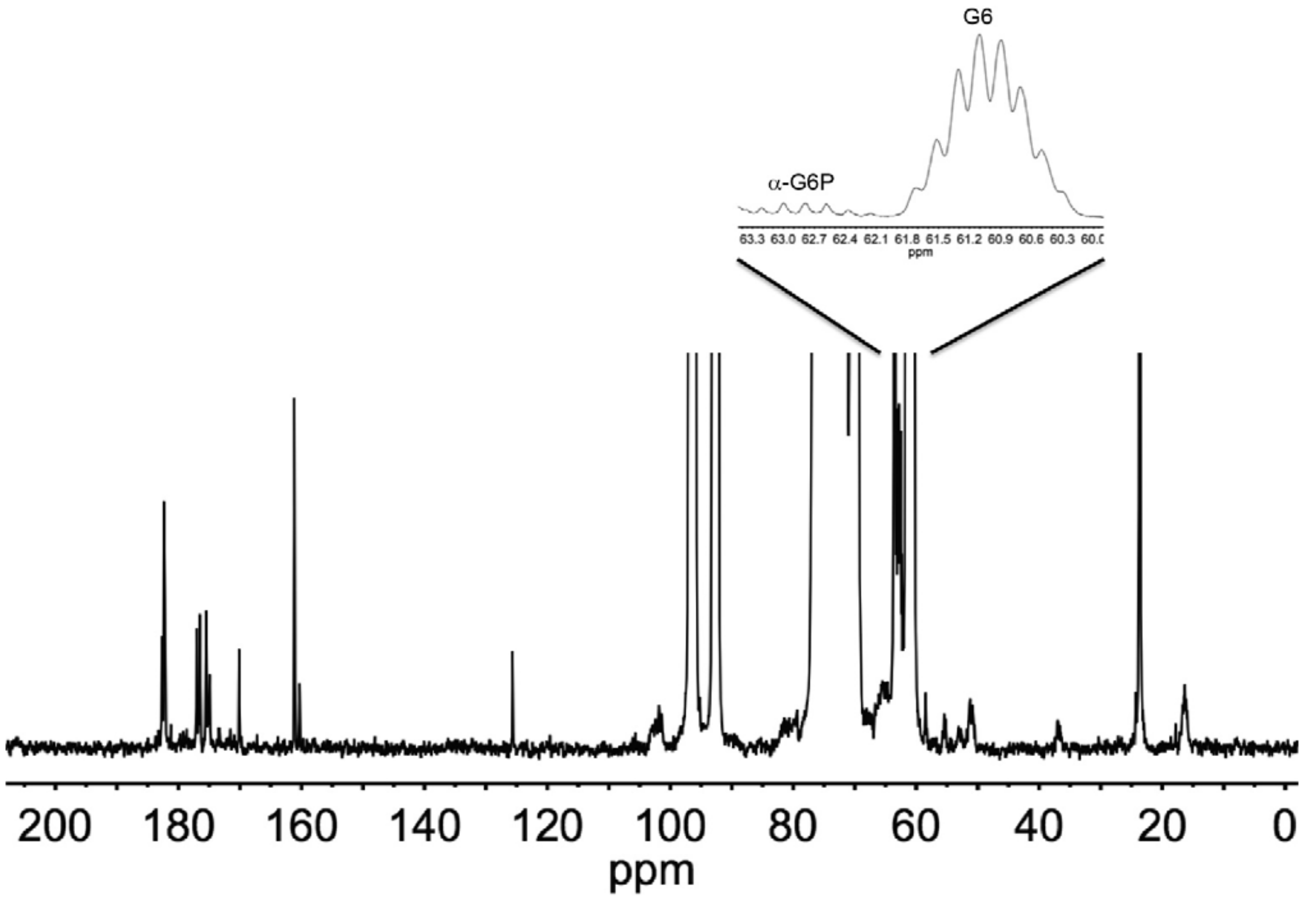


**Figure S5**. *Single $^{13}C$ NMR spectrum of cellular metabolite extract following incubation with isotope labelled glucose for 2 minutes in $D_2O$ buffer*. The insert shows a zoom-in on 6-$^{13}C$-glucose (G6) and the α-form of glucose-6-phosphate (α-G6P).

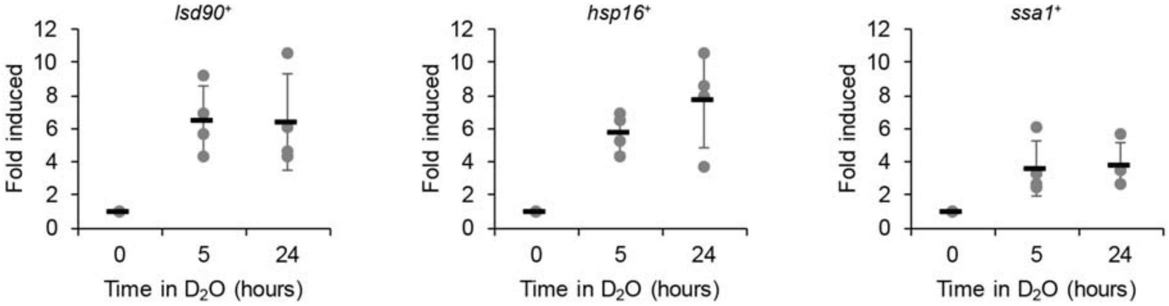


**Figure S6** *Real-time PCR on selected differentially expressed genes*. The amounts of *lsd90*$^+$, *hsp16*$^+$, and *ssa1*$^+$ mRNA were compared by qPCR between cells cultured with 100% $D_2O$ for 0, 5, or 24 hours. Error bars indicate the standard deviation (n=4).

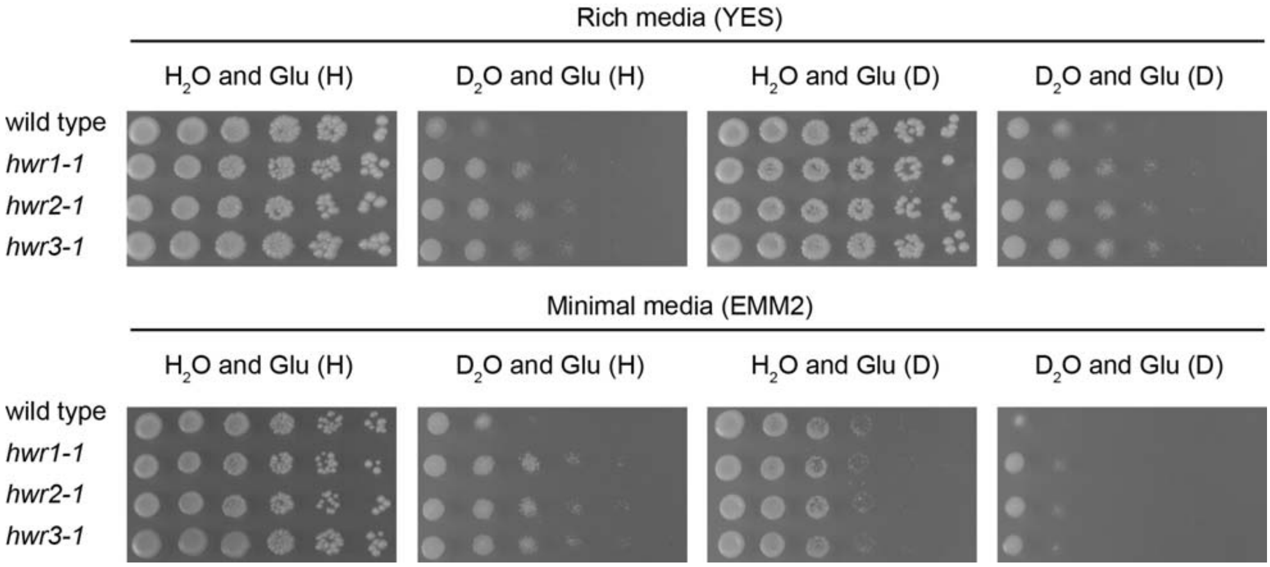


**Figure S7.** *Growth assays of hwr strains on $D_2O$ and deuterated glucose*. The growth of the indicated strains was compared on media prepared with $D_2O$ or $H_2O$ and with either hydrogenated glucose (Glu (H)) or deuterated glucose (Glu (D)) by serial diluting and spotting onto solid rich (YES) media or minimal media (EMM2) agar plates at 30℃.

```
pek1           CTCGCTGGCACATTCACTGGAACTTCGTATTACATGGCGCCTGAACGAATTTCT 756
wt PCR         CTCGCTGGCACATTCACTGGAACTTCGTATTACATGGCGCCTGAACGAATTTCT
hwr1-1 PCR     CTCGCTGGCACATTCACTGGAACTTCGTATTACATGGCGCCTGAACGAATTTCT

pek1           GGGGGATCTTATACTATATCGTCAGATATATGGTCTTTGGGTTTAACATTGATG 810
wt PCR         GGGGGATCTTATACTATATCGTCAGATATATGGTCTTTGGGTTTAACATTGATG
hwr1-1 PCR     GGGGGATCTTATACTATATCGTCAGATATATGGTCTTTGGGTTTAACATTGATG

pek1           GAGGTCGCATTAAACCGGTTTCCATTCCCTCCCGAGGGTAGCCCCCCACCAATG 864
wt PCR         GAGGTCGCATTAAACCGGTTTCCATTCCCTCCCGAGGGTAGCCCCCCACCAATG
hwr1-1 PCR     GAGGTCGCATTAAACCGGTTTCCATTCCCTCCCGAGGGTAGCCCCCCACCAATG

pek1           CCTATTGAATTGCTCTCGTATATAATAAATATGCCACCTCCCCTTTTACCTCAA 918
wt PCR         CCTATTGAATTGCTCTCGTATATAATAAATATGCCACCTCCCCTTTTACCTCAA
hwr1-1 PCR     CCTATTGAATTGCTCTCGTATATAATAAATATGCCACCTCCCCTTTTACCTCAA

pek1           GAACCCGGTATTAAATGGTCGAAATCCTTTCAACATTTTCTATGCGTGTGTCTG 972
wt PCR         GAACCCGGTATTAAATGGTCGAAATCCTTTCAACATTTTCTATGCGTGTGTCTG
hwr1-1 PCR     GAACCCGGTATTAAATGGTCGAAATCCTTTCAACATTTTCTATGCGTGTGTCTG

pek1           GATAAAGACAAAACTCGTCGTCCTGGACCCCAAAAAATGCTTACCCATCCTTGG 1026
wt PCR         GATAAAGACAAAACTCGTCGTCCTGGACCCCAAAAAATGCTTACCCATCCTTGG
hwr1-1 PCR     GATAAAGACAAAACTCGTCGTCCTGGACCCCAAAAAATGCTT-CCCATCCTTGG

pek1           GTTAAAGCGTTTGAGAGAATTCATGTGGACATGGAAGAGTTCCTTCGTCAAGTC 1080
wt PCR         GTTAAAGCGTTTGAGAGAATTCATGTGGACATGGAAGAGTTCCTTCGTCAAGTC
hwr1-1 PCR     GTTAAAGCGTTTGAGAGAATTCATGTGGACATGGAAGAGTTCCTTCGTCAAGTC
```

**Figure S8.** *PCR sequence confirmation of whole genome sequencing on hwr1-1.* Multiple sequence alignment of a segment of the *pek1* gene, and the PCR sequencing results on a wild type (*wt*) strain and the *hwr1-1* strain. Mutation sites are marked in red.

```
mkh1           CAAGGATATAGTGCTAAGGTCGACGTCTGGTCCTTGGGATGTGTAGTGTTGGAA 3078
wt PCR         CAAGGATATAGTGCTAAGGTCGACGTCTGGTCCTTGGGATGTGTAGTGTTGGAA
hwr2-1 PCR     CAAGGATATAGTGCTAAGGTCGACGTCTGGTCCTTGGGATGTGTAGTGTTGGAA

mkh1           ATGTTAGCTGGTCGTAGACCGTGGTCTACAGATGAGGCTATCCAAGCTATGTTC 3132
wt PCR         ATGTTAGCTGGTCGTAGACCGTGGTCTACAGATGAGGCTATCCAAGCTATGTTC
hwr2-1 PCR     ATGTTAGCTGGTCGTAGACCGTGGTCTACAGATGAGGCTATCCAAGCTATGTTC

mkh1           AAG-----------TTAGGTACCGAGAAAAAGGCGCCTCCTATTCCTAGTGAAT 3186
wt PCR         AAG-----------TTAGGTACCGAGAAAAAGGCGCCTCCTATTCCTAGTGAAT
hwr2-1 PCR     AAGCTATGTTCAAGTTAGGTACCGAGAAAAAGGCGCCTCCTATTCCTAGTGAAT

mkh1           TGGTGTCTCAGGTATCACCCGAAGCGATTCAATTTTTGAATGCATGCTTTACTG 3240
wt PCR         TGGTGTCTCAGGTATCACCCGAAGCGATTCAATTTTTGAATGCATGCTTTACTG
hwr2-1 PCR     TGGTGTCTCAGGTATCACCCGAAGCGATTCAATTTTTGAATGCATGCTTTACTG

mkh1           TGAATGCTGATGTAAGGCCAACCGCAGAGGAATTATTAAATCACCCGTTTATGA 3294
wt PCR         TGAATGCTGATGTAAGGCCAACCGCAGAGGAATTATTAAATCACCCGTTTATGA
hwr2-1 PCR     TGAATGCTGATGTAAGGCCAACCGCAGAGGAATTATTAAATCACCCGTTTATGA
```

**Figure S9.** *PCR sequence confirmation of whole genome sequencing on hwr2-1.* Multiple sequence alignment of a segment of the *mkh1* gene, and the PCR sequencing results on a wild type (*wt*) strain and the *hwr2-1* strain. Mutation sites are marked in red.

```
pck2           CCTGAATTTATGGCACCGGAAATCTTATTAGAACAGCAATATACGAGAAGCGTT 2592
wt PCR         CCTGAATTTATGGCACCGGAAATCTTATTAGAACAGCAATATACGAGAAGCGTT
hwr3-1 PCR     CCTGAATTTATGGCACCGGAAATCTTATTAGAACAGCAATATACGAGAAGCGTT

pck2           GACTGGTGGGCTTTTGGTGTACTAATTTACCAAATGCTGCTTGGTCAATCTCCA 2646
wt PCR         GACTGGTGGGCTTTTGGTGTACTAATTTACCAAATGCTGCTTGGTCAATCTCCA
hwr3-1 PCR     GACTGGTGGGCTTTTAGTGTACTAATTTACCAAATGCTGCTTGGTCAATCTCCA

pck2           TTTAGAGGAGAAGACGAAGAAGAAATTTTTGATGCAATTTTATCTGATGAACCT 2700
wt PCR         TTTAGAGGAGAAGACGAAGAAGAAATTTTTGATGCAATTTTATCTGATGAACCT
hwr3-1 PCR     TTTAGAGGAGAAGACGAAGAAGAAATTTTTGATGCAATTTTATCTGATGAACCT

pck2           TTGTATCCTATTCATATGCCAAGGGATTCCGTTTCTATATTACAACAACTTTTG 2754
wt PCR         TTGTATCCTATTCATATGCCAAGGGATTCCGTTTCTATATTACAACAACTTTTG
hwr3-1 PCR     TTGTATCCTATTCATATGCCAAGGGATTCCGTTTCTATATTACAACAACTTTTG

pck2           ACTCGCGATCCTAAAAAACGACTTGGATCTGGCCCTAACGATGCAGAAGATGTC 2808
wt PCR         ACTCGCGATCCTAAAAAACGACTTGGATCTGGCCCTAACGATGCAGAAGATGTC
hwr3-1 PCR     ACTCGCGATCCTAAAAAACGACTTGGATCTGGCCCTAACGATGCAGAAGATGTC
```

**Figure S10.** *PCR sequence confirmation of whole genome sequencing on hwr3-1.* Multiple sequence alignment of a segment of the *pck2* gene, and the PCR sequencing results on a wild type (*wt*) strain and the *hwr3-1* strain. Mutation sites are marked in red.

**Table S1**

*Differentially expressed tRNAs after 24 hours in $D_2O$*

| Systematic ID | tRNA | Expression change (fold) | FDR* |
|---|---|---|---|
| *SPATRNASER.01* | Ser, cytosolic | 0.14 | 0.0087 |
| *SPATRNASER.03* | Ser, cytosolic | 0.16 | 0.0087 |
| *SPATRNAMET.01* | Met, cytosolic | 0.16 | 0.0087 |
| *SPMITTRNALEU.01* | Leu, mitochondrial | 0.17 | 0.0087 |
| *SPMITTRNAPHE.01* | Phe, mitochondrial | 0.18 | 0.0087 |
| *SPMITTRNAVAL.01* | Val, mitochondrial | 0.18 | 0.0087 |
| *SPMITTRNAARG.02* | Arg, mitochondrial | 0.18 | 0.0087 |
| *SPMITTRNAGLY.01* | Gly, mitochondrial | 0.19 | 0.0087 |
| *SPMITTRNAMET.02* | Met, mitochondrial | 0.20 | 0.0087 |
| *SPMITTRNATYR.01* | Tyr, mitochondrial | 0.20 | 0.0087 |
| *SPMITTRNAILE.02* | Ile, mitochondrial | 0.21 | 0.0087 |
| *SPCTRNAMET.07* | Met, cytosolic | 0.21 | 0.0087 |
| *SPMITTRNAHIS.01* | His, mitochondrial | 0.22 | 0.0087 |
| *SPMITTRNAPRO.01* | Pro, mitochondrial | 0.23 | 0.0087 |
| *SPMITTRNASER.02* | Ser, mitochondrial | 0.23 | 0.0087 |
| *SPATRNAMET.03* | Met, cytosolic | 0.24 | 0.0087 |
| *SPMITTRNAASP.01* | Asp, mitochondrial | 0.24 | 0.0087 |
| *SPMITTRNALEU.02* | Leu, mitochondrial | 0.25 | 0.0087 |
| *SPBTRNAMET.05* | Met, cytosolic | 0.25 | 0.0087 |
| *SPMITTRNACYS.01* | Cys, mitochondrial | 0.25 | 0.0087 |
| *SPMITTRNATRP.01* | Trp, mitochondrial | 0.25 | 0.0087 |
| *SPMITTRNAGLN.01* | Gln, mitochondrial | 0.26 | 0.0087 |
| *SPATRNATYR.01* | Tyr, cytosolic | 0.34 | 0.0203 |
| *SPBTRNALYS.07* | Lys, cytosolic | 0.37 | 0.0267 |
| *SPMITTRNAARG.01* | Arg, mitochondrial | 0.39 | 0.0087 |
| *SPATRNAPRO.02* | Pro, cytosolic | 0.40 | 0.0092 |
| *SPMITTRNAILE.01* | Ile, mitochondrial | 0.42 | 0.0088 |
| *SPMITTRNAALA.01* | Ala, mitochondrial | 0.43 | 0.0087 |
| *SPATRNALEU.01* | Leu, cytosolic | 0.44 | 0.0393 |
| *SPATRNALEU.03* | Leu, cytosolic | 5.28 | 0.0087 |

*False discovery rate.

**Table S2**

*Amino acid changes in affected kinases in the hwr strains*

| Mutant | Protein | aa change |
|---|---|---|
| *hwr1-1* | Pek1 | L338LPILGLKRLREFMWTWKSSFVKSGLIRRNSVICstop |
| *hwr2-1* | Mkh1 | L1046LCSSstop |
| *hwr3-1* | Pck2 | G870S |
